# Homer condensates orchestrate YAP-Wnt signaling crosstalk downstream of the Crumbs polarity complex

**DOI:** 10.1101/2025.07.23.666266

**Authors:** Siti Maryam J M Yatim, Linda Jiabao Woo, Yuhong Chen, Barbara Hübner, Alexander Ludwig

## Abstract

The Hippo pathway governs cell growth, proliferation, and differentiation and is frequently deregulated in cancer. YAP, the central transcriptional co-activator of the Hippo pathway, is suppressed by diverse upstream signals including cell density and polarity. YAP also functionally interacts with the Wnt/β-catenin pathway, yet how polarity cues coordinate this pathway crosstalk remains poorly understood. Here, we demonstrate that Homer scaffolding proteins couple the Crumbs polarity complex to the coordinated regulation of YAP and Wnt signaling. Homers directly interact, via their EVH1 domains, with the Crumbs component PATJ and the NDR kinase scaffold Furry-like (FRYL). Functionally, Homers antagonize FRYL to promote YAP activation while cooperating with FRYL to enhance Wnt/β-catenin signaling, revealing pathway-selective regulation. PATJ, in contrast, acts upstream to restrain Homer-driven YAP signaling. Interestingly, in non-polarized epithelial and colorectal cancer cells, Homers form cytoplasmic biomolecular condensates whose assembly and material properties are differentially modulated by PATJ and FRYL. Whereas FRYL promotes spherical, liquid-like Homer condensates, PATJ drives the formation of irregular, network-like assemblies, thereby altering condensate topology and signaling output. Collectively, our findings establish Homer-driven phase separation as a tunable signaling mechanism that integrates polarity cues with YAP–Wnt pathway coordination and transcriptional output.

**Significance:** The Hippo/YAP and Wnt signaling pathways play important roles in development and are frequently deregulated in cancer. Both pathways respond to epithelial cell architecture, but how polarity cues coordinate their crosstalk remains poorly understood. Here, we identify Homer proteins as phase-separating scaffolds that link apico-basal polarity to YAP-Wnt pathway integration. We show that the material properties of Homer condensates are tunable and depend on protein abundance and binding partners, providing a mechanism by which polarity cues can shape signaling output through changes in phase behavior. Our findings establish Homer condensates as a polarity-sensitive signaling hub that modulates YAP transcriptional programs and coordinates YAP-Wnt pathway crosstalk.

## Introduction

The Hippo pathway is a central regulator of cell growth, proliferation, differentiation, and survival, and is frequently deregulated in cancer. Its core effectors are the transcriptional coactivators Yes-associated protein (YAP) and its paralogue TAZ (WWTR1), which shuttle between the cytoplasm and nucleus to control gene expression. In the canonical pathway, the Ste20-like kinases MST1/2 activate the Large Tumor Suppressor kinases LATS1/2, which phosphorylate YAP/TAZ on multiple serine/threonine residues. This promotes 14-3-3 binding, cytoplasmic retention, and proteasomal degradation, thereby preventing YAP/TAZ from activating TEA domain transcription factors (TEADs) in the nucleus (1, 2). In addition to this core kinase cassette, several non-canonical kinases—including MST3/4, NDR1/2, MAP4Ks, and MEKK family members—also modulate YAP activity (2–6). Recent studies further indicate that Hippo pathway components can undergo phase separation, suggesting that biomolecular condensates contribute to pathway regulation. These assemblies appear responsive to diverse upstream inputs, including osmotic stress, actin cytoskeletal tension, extracellular matrix stiffness, cell–cell junctions, and polarity cues (7–14). However, how such signals control the signaling output of phase separated Hippo regulators is poorly understood.

YAP is closely intertwined with other signaling pathways including the TGFβ, NOTCH, and Wnt pathways (15–19). Several studies have demonstrated that YAP/TAZ play a dual role in regulating Wnt/β-catenin signaling (20–24). In the absence of a Wnt signal, cytoplasmic YAP/TAZ are incorporated into the β-catenin destruction complex via an interaction with Axin1 to promote β-catenin degradation (21). However, when Wnt signaling is activated, YAP/TAZ dissociate from the destruction complex and translocate into the nucleus to form transcriptional complexes with β-catenin and the TCF/LEF transcription factors, leading to the expression of YAP/TAZ and Wnt-dependent target genes (21, 24). YAP/TAZ may also suppress Wnt signaling by binding to Disheveled, which prevents its activation (20), or by binding directly to β-catenin, which stabilizes β-catenin in the cytoplasm (23). Moreover, both canonical and non-canonical Wnt signaling can feedback to the Hippo pathway to control the activity of YAP (25, 26). Such complex reciprocal interactions between the YAP/TAZ and Wnt pathways have significant implications during differentiation, regeneration, and in cancers (27–29), yet how this pathway crosstalk is regulated remains largely unclear.

The epithelial Crumbs complex, which is composed of the transmembrane protein Crumbs (Crb), the adaptor protein Pals1, and the multi-PDZ domain scaffolding protein PATJ, is an evolutionarily conserved regulator of cell polarity and an important upstream regulator of the Hippo pathway (30–34). In *Drosophila*, Crb restricts tissue growth by promoting apical retention and inhibition of Yorkie (the YAP/TAZ orthologue) (35–38). Similarly, loss of Crb3 or Pals1 in mammalian cells and mouse models leads to YAP/TAZ hyperactivation, accompanied by defects in tissue architecture and differentiation (39–42). Despite these observations, the molecular mechanisms linking the Crumbs complex to Hippo/YAP regulation—particularly in mammalian epithelial cells—remain incompletely defined.

We previously demonstrated that the mammalian Crumbs complex defines a distinct cortical domain apical of epithelial tight junctions, termed the vertebrate marginal zone (VMZ) (32, 43). In that study we identified the Homer scaffolding proteins as novel PATJ interactors. Mammalian Homers (Homer1-3) are composed of an N-terminal EVH1 domain that binds PPxxF motif–containing ligands and a C-terminal coiled coil domain that mediates homo- and hetero-tetramerization (44–46). This tetrameric organization confers multivalency, a key feature underlying their ability to assemble biomolecular condensates with postsynaptic density proteins (47–49). Beyond their established roles in synaptic signaling and cytoskeletal organization, emerging evidence implicates Homers in calcium-dependent mechanosensing at epithelial cell junctions (50), in neutrophil cell polarity and migration (51), and in YAP and β-catenin signaling in cancer cells (52–56), suggesting that they may link polarity cues to growth control pathways.

Here, using complementary loss- and gain-of-function approaches combined with transcriptomics, qPCR, and reporter assays, we demonstrate that Homers enhance YAP/TEAD and canonical Wnt signaling in MDCK, 293T, and HCT116 colorectal cancer cells. Homers promote YAP/TEAD signaling through the assembly of cytoplasmic biomolecular condensates, whose assembly, material properties, and signaling functions are differentially modulated by PATJ and the NDR kinase scaffolding protein FRYL. Collectively, our data indicate that Homers orchestrate YAP-Wnt signaling crosstalk in cancer and non-cancer cells through an adaptable phase separation-dependent signaling mechanism.

## Results

### PATJ directly interacts with and recruits Homers to apical cell junctions

Homers are abundantly expressed in the brain but are also detected in various non-neuronal tissues including the heart, lung, intestine, and kidney (57). Using GFP-tagged Homer proteins we previously demonstrated that all Homers localize to the apical-lateral border in fully polarized Madin Darby Canine Kidney type II (MDCK-II) cells (43). To verify this, we performed immunofluorescence confocal microscopy of MDCK monolayers using specific antibodies against Homer1, 2, and 3 and the tight junction marker ZO-1. All endogenous Homers localized to apical cell junctions, as expected (Fig. 1A and Fig. S1A). Moreover, antibodies against Homer1, which is the only Homer protein expressed in the kidney (57), produced an apical junctional and punctate cytoplasmic staining in mouse kidney sections that partially colocalized with ZO-1 (Fig. S1B-S1D) (Pearson’s r = 0.60 ± 0.08; n=30). We concluded that Homers are integral components of apical cell junctions in renal epithelial cells.

**Figure 1:**
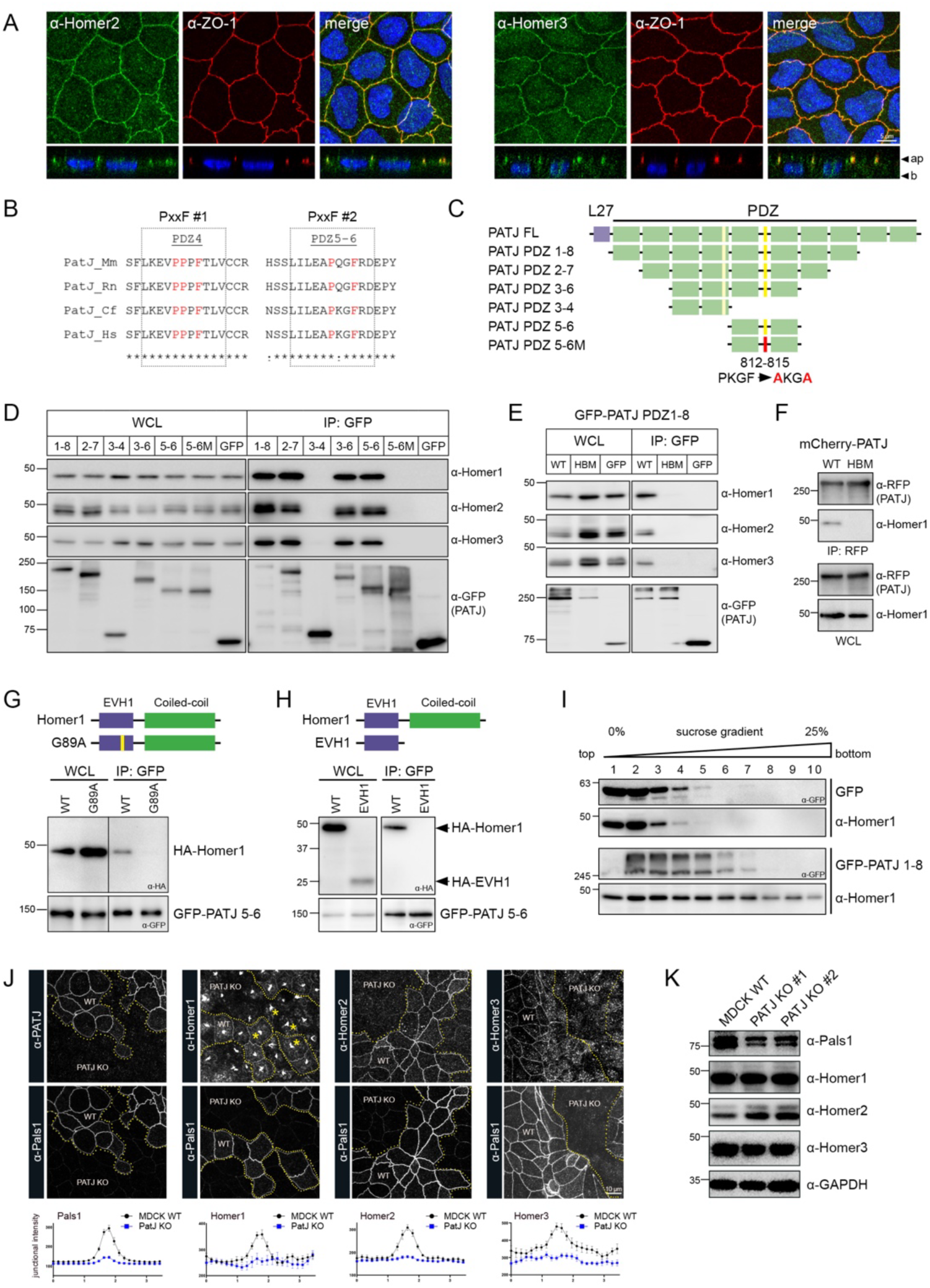
Homers are recruited to apical cell junctions via a direct interaction with PATJ. (A) Airyscan microscopy of endogenous Homers in MDCK-II cells. Cells were grown to confluency on Transwell filters for 14 days, fixed and co-stained with ZO-1 and Homer2 or Homer3 antibodies. XY (en face) and XZ (side view) projections are shown. (B) Amino acid sequence alignment of the two conserved PxxF motifs in PATJ. Mm: *Mus musculus*, Rn: *Rattus norvegicus*, Cf: *Canis lupus familiaris*, Hs: *Homo sapiens* (C) Schematic representation of PATJ PDZ domain constructs cloned into the APEX2-EGFP-C1 (A2E-C1) expression vector. (D) Co-immunoprecipitation (Co-IP) assay in 293T cells transfected with the constructs shown in (C). The interaction with endogenous Homer proteins was analysed by Western blotting (WB). (E) Co-IP assay of GFP-PATJ PDZ1-8 and the corresponding PxxF mutant. (F) Co-IP assay of full-length wild-type mCherry-PATJ and the corresponding PxxF mutant. (G) Co-IP assay of GFP-PATJ PDZ5-6 with wild-type HA-Homer1 or the EVH1 domain mutant (G89A). (H) Co-IP assay of GFP-PATJ PDZ5-6 with HA-Homer1 or the HA-tagged Homer1 EVH1 domain. (I) Sucrose density gradient centrifugation of 293T cell lysates. Cells were transfected with the A2E-C1 construct (control; top) or the A2E-PATJ PDZ1-8 construct (bottom). Gradient fractions were probed with GFP and Homer1 antibodies. (J) Maximum intensity projections of confocal z-stacks of MDCK wild-type (WT) and PATJ KO cells co-cultured on Transwell filters for 12 days. Pals1 antibody served as a marker for WT cells. Note that anti-Homer1 antibodies produce a non-specific staining of unknown origin (yellow asterisks). Bottom: Average junctional signal intensity of Pals1 and Homers in PATJ KO and MDCK WT cells determined by line scan analysis (n=11 each). (K) WB analysis of MDCK WT and PATJ KO MDCK cell lysates probed for Pals1, PATJ and Homers.

We previously demonstrated an interaction between PATJ and Homers using co-immunoprecipitations (co-IPs) (43). PATJ is a large PDZ-domain containing protein that interacts with the Hippo pathway regulators angiomotin (AMOT) and KIBRA (32, 58–60) and to the tight junction proteins claudin-1 and ZO-1 or ZO-3 (32, 61, 62). To define the molecular basis of the Homer/PATJ interaction, we screened PATJ for PPxxF motifs that might provide docking sites for the Homer EVH1 domains. Two highly conserved PxxF motifs were identified in PATJ: One (PPPF) at the C-terminus of the PDZ4 domain, and another (PKGF) in the linker between PDZ5 and PDZ6 (Fig. 1B). To test whether these motifs are required for the interaction with Homers, GFP-tagged PATJ PDZ domain fragments were expressed in 293T cells and immunoprecipitated, followed by Western blotting for endogenous Homers (Fig. 1C). A fragment containing PDZ5-6 was sufficient to interact with Homers and mutation of the PKGF motif abolished the binding (Fig. 1D). Inactivation of the PKGF Homer binding site in the PATJ PDZ1-8 fragment (Fig. 1E) or in full-length PATJ (containing the L27 domain) (Fig. 1F) also eliminated the interaction, demonstrating that this sequence motif is critical for Homer association. Moreover, mutation of glycine 89 (G89) in the Homer1 EVH1 domain, which is essential for ligand binding (63), abolished the interaction with PATJ (Fig. 1G) and interfered with the recruitment of Homers to cell-cell junctions in MDCK cells (not shown). Interestingly, the Homer1 EVH1 domain alone was insufficient to interact with PATJ (Fig. 1H), suggesting that Homer oligomerization via the coiled-coil domains is required for a stable interaction with PATJ. To test whether PATJ oligomerizes with Homers, we performed sucrose density gradient centrifugation. Indeed, GFP-PATJ PDZ1-8 partially co-fractionated with Homer1 in the gradient and markedly increased the size or density of Homer oligomers compared to control-transfected cells (Fig. 1I). We concluded that PATJ interacts directly with the EVH1 domain of Homers via a conserved PxxF motif located in the linker between PDZ domains 5 and 6 to assemble higher order hetero-oligomers.

To test whether PATJ is required for the recruitment of Homers to the apical-lateral border, we co-cultured MDCK wild-type (WT) and PATJ knockout (KO) cells on Transwell filters and stained such co-cultures with antibodies against each Homer protein. Antibodies against Pals1, which is lost from apical cell junctions in the absence of PATJ (64), were used to discriminate between WT and PATJ KO cells. In WT cells, all Homers localized to the apical-lateral border, as expected. By contrast, in cells lacking PATJ, all Homer proteins were clearly displaced from apical cell junctions (Fig. 1J). Western blotting showed that the expression of Homers was not notably altered in PATJ KO cells (Fig. 1K). Hence, PATJ is essential for the recruitment of Homers to apical cell junctions in polarized epithelia.

### Homers and PATJ coordinate YAP/TEAD and Wnt signaling in polarised epithelial cells

To address the functions of individual Homers and the entire Homer protein family, we generated single and triple Homer KO MDCK cells using CRISPR/Cas9 gene editing. All KO lines were validated by PCR/DNA sequencing of the guide RNA target locus, Western blotting, and immunofluorescence (Fig. S2A-S2F). Cortical actin levels were slightly increased in Homer triple KO (TKO) cells, whereas the localization of PATJ and Pals1 and several junctional proteins was not affected (Fig. S2G and S2H). This indicates that Homers are dispensable for the assembly of the Crumbs complex and apical cell junctions. Homer TKO cells also showed no obvious defects in cell proliferation (now shown). However, scratch wound assays of confluent MDCK cell monolayers indicated that Homers are required for efficient wound closure (Fig. S2I), suggesting a function in collective cell migration.

Next, we analysed the transcriptome of Homer TKO cells using RNA sequencing (Fig. S3A, Table S2). Interestingly, genes involved in cancer signaling were significantly deregulated in Homer TKO cells (Fig. 2A, 2B and Fig. S3B). Closer inspection of the differentially expressed genes (DEGs) showed that the canonical YAP/TEAD target genes CTGF, ANKRD1 and GADD45A (65) were downregulated in Homer TKO cells (Fig. 2C). Other potential YAP/TEAD target genes including FOLR2 (66), COL12A1 (67, 68), and HOXA2 (69) were downregulated as well, and so was RASSF1, an important regulator of the Hippo pathway. In addition, the expression of several genes involved in Wnt signaling was reduced in Homer TKO cells (Fig. 2C). This includes the Wnt receptor FZD4, the Wnt target genes HAS2 (70–73), ARHGAP24 (74), and POU3F3 (75), and GAS1 (76), DACH1 (77–79), PLEKHA4 (80), and GNA14 (81). To verify the transcriptional changes observed by RNA sequencing, we performed qPCRs. In addition, TEAD (8xGTIIC) and TCF/LEF (TOPFlash) luciferase reporter assays were used to analyse Hippo/YAP and canonical Wnt/β-catenin pathway activity, respectively. As expected, TEAD reporter activity and the expression of CTGF and ANKRD1 was significantly reduced in Homer TKO cells (Fig. 2E and 2F). Homer TKO cells also showed a 1.2-1.5-fold increase in YAP S127 phosphorylation (Fig. 2D), indicating a suppression of YAP activation. qPCR analysis further confirmed that FZD4, GNA14, DACH1, and HAS2 were downregulated in Homer TKO cells (Fig. 2G). In addition, although the expression of the canonical Wnt target gene AXIN2 was unchanged (Fig. 2H), TOPFlash reporter activity was markedly reduced in Homer TKO cells (Fig. 2I).

**Figure 2:**
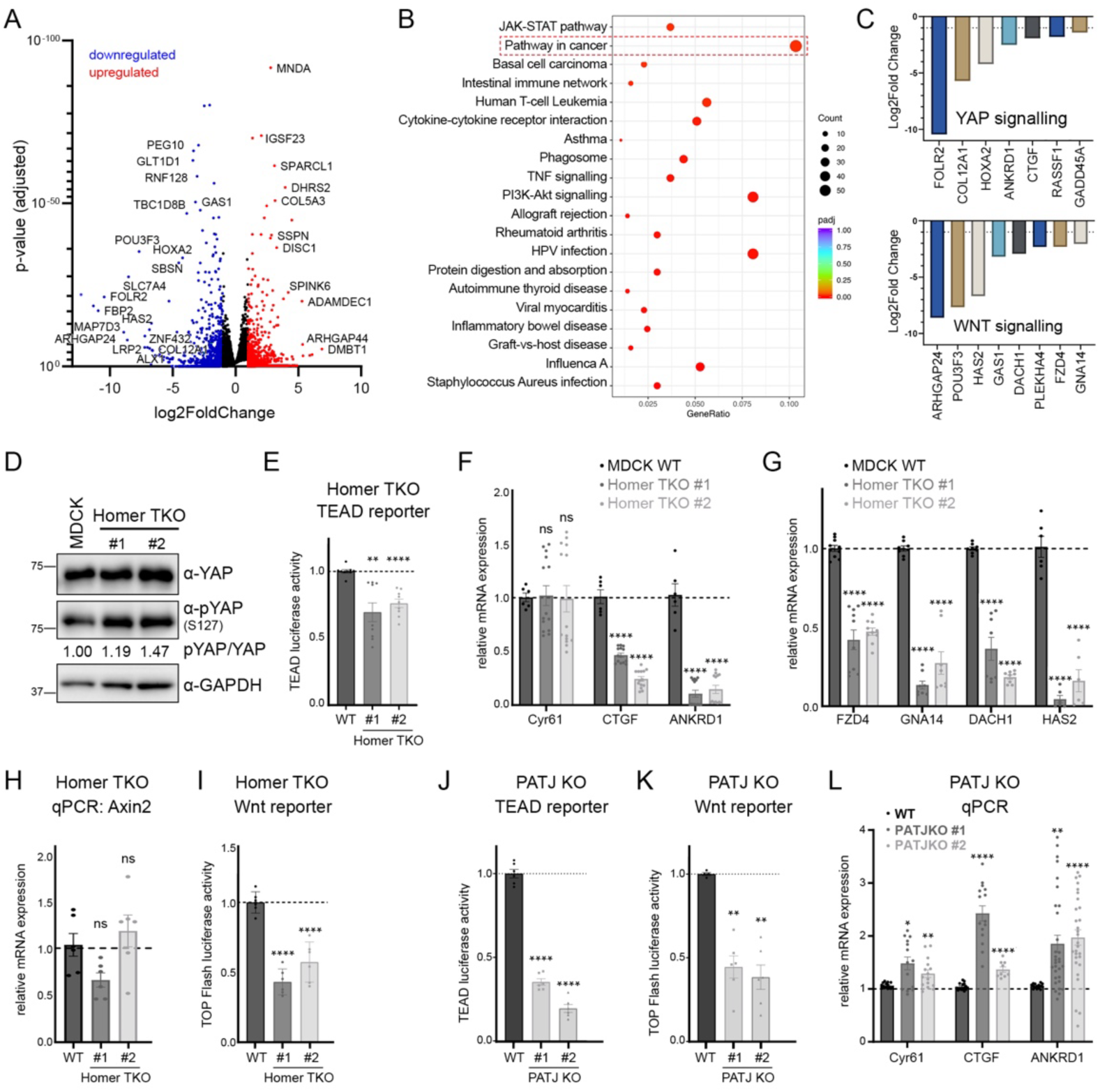
Homers and PATJ functionally interact to regulate YAP/TEAD and Wnt pathway activity. (A) Volcano plot showing select differentially expressed genes (DEGs) in Homer TKO cells identified by RNA-sequencing. (B) KEGG pathway analysis of DEGs in Homer TKO cells. The y-axis lists significantly enriched pathways, while the x-axis represents the gene ratio. Dot size indicates the number of DEGs involved in each pathway, and colour reflects the adjusted p-value (padj), with red indicating higher statistical significance. (C) Bar graphs showing the log2 fold changes of DEGs associated with the YAP (top) and WNT (bottom) signalling pathways. Genes were selected based on their relevance to each pathway. (D) WB analysis of WT MDCK and Homer TKO cell lysates probed for YAP and pYAP (S127). The pYAP/YAP ratio was quantified by densitometry (n=3). (E) TEAD luciferase activity in Homer TKO cells (n=4). (F and G) qPCR analysis of canonical YAP target genes (F) and WNT-associated genes (G) in Homer TKO cells (n=4). (H) qPCR analysis of Axin2 expression in Homer TKO cells (n=4). (I) TOPFlash luciferase activity in Homer TKO cells (n=3). (J and K) TEAD and TOPFlash luciferase reporter assays in PATJ KO cells (n=3). (L) qPCR analysis of YAP target genes in PATJ KO cells (n=6). Luciferase assay data were analysed using a paired Student’s t-test. qPCR data were analysed using Two-Way ANOVA. Data is presented as mean ± SEM. * P ≤ 0.05, ** P ≤ 0.01, *** P ≤ 0.001, **** P ≤ 0.0001.

Our data so far indicated that Homers promote YAP/TEAD and canonical Wnt signaling. Loss of PATJ, which resulted in the removal of Homers from apical cell junctions (Fig. 1J), would therefore be expected to equally alter the signaling output of these pathways. Indeed, both TEAD and Wnt reporter activities were significantly reduced in PATJ KO cells (Fig. 2J and 2K). Moreover, YAP target gene transcription was increased (Fig. 2L), while HAS2 and FZD4 mRNA levels were decreased (not shown). We concluded that Homers link the Crumbs complex to YAP/TEAD and Wnt pathway regulation, and that PATJ functionally antagonizes Homer-driven YAP target gene expression through a TEAD-independent mechanism.

To understand whether Homers function synergistically or in a unique fashion, we examined single Homer KO cells. Interestingly, whereas loss of *Homer3* reduced YAP/TEAD signaling and increased YAP S127 phosphorylation, inactivation of *Homer1* or *Homer2* unexpectedly enhanced YAP activation. Consistent with this, overexpression of Homer3—but not Homer1—stimulated both TEAD and TOPFlash reporter activity (Fig. S3C-3H). This indicates that the three Homer paralogues are not functionally redundant. Although combined loss of all Homers dampens YAP/TEAD and Wnt signaling, individual Homers exert distinct and, in part, opposing effects. We concluded that Homer3 acts as the dominant pro-YAP/Wnt regulator in MDCK cells, and that the overall pathway output reflects the balance between paralogue-specific activities.

### Homers promote YAP and Wnt pathway activity in 293T and HCT116 cells

We next analysed the functions of Homers in cells lacking polarity cues. In contrast to polarized MDCK cells, siRNA-mediated silencing of individual or all three Homers in 293T cells uniformly reduced YAP target gene expression and TOPFlash reporter activity, whereas overexpression of Homers was associated with elevated TEAD and TOPFlash reporter readouts (Fig. S4A-S4E). Notably, TEAD activation was independent of Homer-mediated actin bundling (47), as wild-type Homer3 and an actin-binding–deficient mutant stimulated the reporter to a similar extent (Fig. S4F). Together, these results indicated that Homers promote YAP and canonical Wnt signaling in both polarized and non-polarized epithelial cells, and that their paralogue-specific functions are cell-type or context-dependent.

Given the prominent role of YAP-Wnt pathway crosstalk in colorectal cancer (CRC) (22, 23, 25, 82), we next assessed Homer function in HCT116 cells, which harbor oncogenic KRAS and a stabilizing 3-bp deletion in *CTNNB1* that enhances β-catenin-dependent Wnt signaling (83). HCT116 cells expressed Homer1 and Homer3, but not Homer2 (Fig. S4G). siRNA-mediated Homer depletion significantly reduced YAP/TEAD and TOPFlash reporter activity without markedly altering the pYAP/YAP ratio or total β-catenin levels (Fig. S4H-S4J). Importantly, the expression of canonical YAP (CTGF, PTPN14) and Wnt target genes (HAS2, SOX9, AXIN2) was significantly decreased in cells treated with Homer siRNA (Fig. S4K and S4L). Moreover, and in agreement with our MDCK cell data, Homer knockdown did not affect cell proliferation (not shown), but markedly impaired wound closure in scratch wound assays (Fig. S4M). Together, these findings identify Homers as positive regulators of YAP and Wnt signaling in CRC cells and suggest a role in promoting cancer cell migration.

### Homers control YAP/TEAD signaling via the NDR kinase scaffolding protein FRYL

To identify proteins that could link Homers to the YAP and Wnt signaling pathways, we performed immunoprecipitation-mass spectrometry (IP-MS) of GFP tagged Homer3 stably expressed in MDCK cells (Fig. S5A). As expected, Homer1 and Homer2 as well as PATJ were present in Homer3 IPs. We further identified the Furry-like protein (FRYL) as a novel Homer3 interacting protein. Furry, the FRYL paralogue, is a conserved scaffolding protein of NDR1/2 kinases (Tricornered in *Drosophila*). Both Furry and NDR1/2 have been shown to regulate YAP (4, 5) and are critical for vertebrate (84) and invertebrate (85–89) development. To ascertain that FRYL interacts with NDR, as suggested previously (90) (Fig. 3A), we performed IP experiments in 293T cells. GFP-NDR1 co-immunoprecipitated with FRYL and the NDR scaffolding protein MOB1a, whereas LATS1 was not detected in GFP-NDR1 IPs (Fig. 3B). In addition, overexpression of FRYL increased NDR Thr442/Thr444 phosphorylation (a readout for NDR kinase activity) but did not affect the phosphorylation of LATS1/2 (Thr1029) (Fig. 3C). This establishes FRYL as a specific NDR1 scaffolding protein.

**Figure 3:**
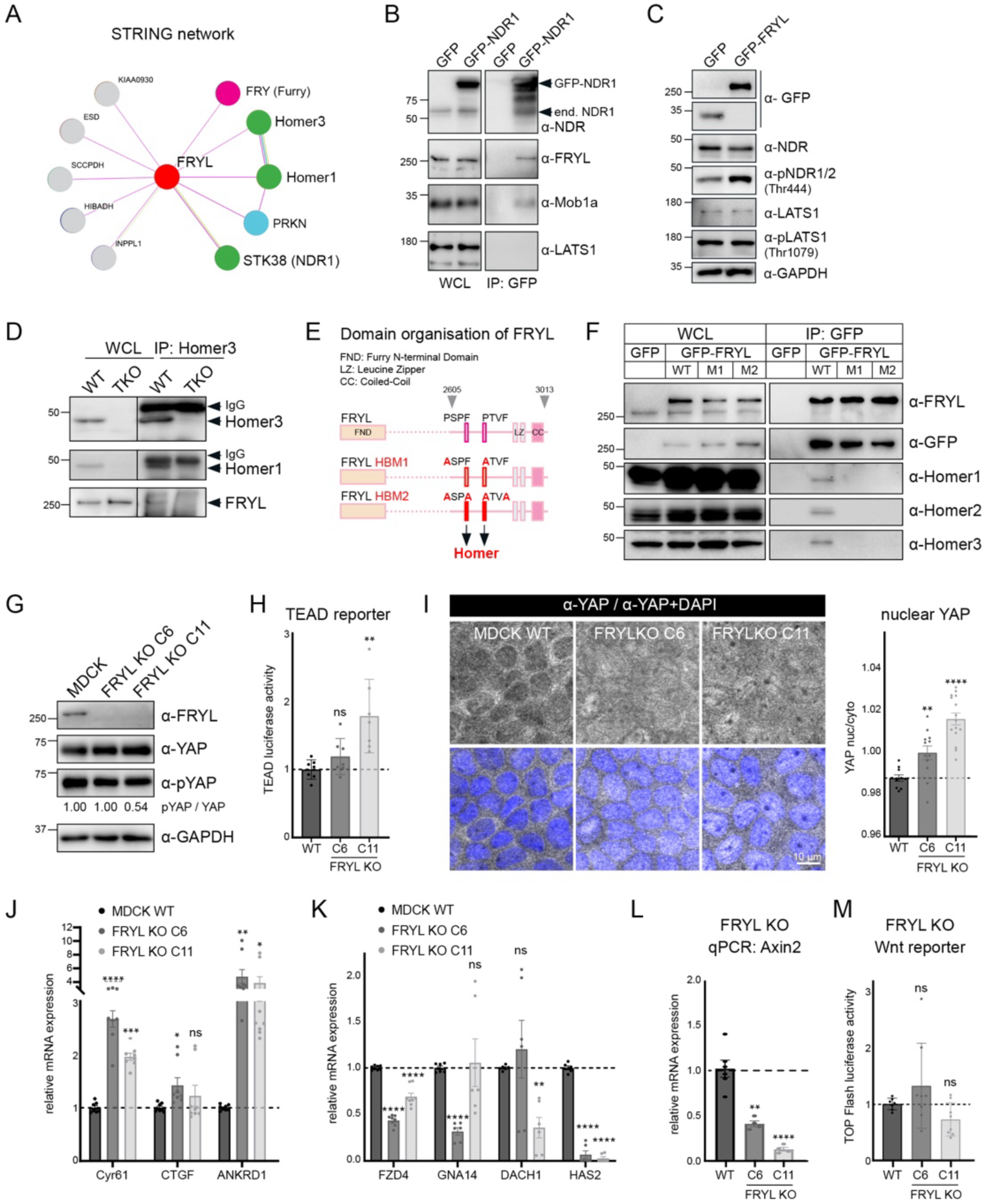
The NDR kinase scaffolding protein FRYL interacts with Homers and suppresses YAP/TEAD signaling. (A) STRING interactome of FRYL based on evidence for physical interactions. (B) IP of GFP-NDR1 from 293T cell lysates. (C) WB analysis of 293T cells transfected with GFP or GFP-FRYL. (D) IP of the endogenous Homer/FRYL complex from MDCK cell lysates using anti-Homer3 antibodies. Lysates from Homer TKO cells were used as a control. (E) Domain organisation of FRYL. The two Homer binding sites are depicted. The Homer binding mutants (HBM) M1 and M2 exhibit distinct mutations in the two PxxF motifs. (F) IP of WT GFP-FRYL and the two FRYL HBMs M1 and M2 from 293T cell lysates. (G) WB analysis of MDCK WT and FRYL KO cell lysates probed for FRYL, YAP and p-YAP (S127). The p-YAP/YAP ratio was quantified by densitometry (n=3). (H) TEAD luciferase activity in FRYL KO cells normalised to MDCK WT cells (n=3). (I) FRYL KO and MDCK WT cells were cultured on Transwell filters, fixed and stained for total YAP. Nuclear and cytoplasmic YAP intensities were quantified using DAPI staining as a mask to define the nucleus. Quantification was performed on 10 images per condition, analyzing more than 80 cells per image (n = ∼800 cells). Data is presented as mean ± SEM. **P ≤ 0.01, ****P ≤ 0.0001 (Student’s t-test). (J and K) qPCR analysis of YAP target genes (J) and WNT-associated genes (K) in FRYL KO cells (n=3). (L) qPCR analysis of Axin2 mRNA levels in FRYL KO cells (n=3). (M) TOPFlash luciferase activity in FRYL KO cells normalized to MDCK WT cells (n=3). All Luciferase assays were statistically analysed using a paired Student’s t-test. All qPCR data were analysed using Two-Way ANOVA Tukey multiple comparison test. Data is presented as mean ± SEM. *P ≤ 0.05; **P ≤ 0.01; ***P ≤ 0.001; ****P ≤ 0.0001.

To confirm that Homers interact with FRYL, we immunoprecipitated endogenous Homer3 from MDCK cell lysates. As expected, Homer1 and FRYL were present in Homer3 IPs, indicating that Homer oligomers form a complex with FRYL in polarized epithelia (Fig. 3D). We noted that the C-terminus of FRYL harbors two highly conserved PxxF motifs that could potentially function as Homer binding sites (Fig. 3E and Fig. S5B-S5D). Indeed, wild-type GFP-FRYL efficiently interacted with all three Homers in IPs, whereas no interactions were observed with two independent Homer binding mutants (HBMs) in which the two PxxF motifs were mutated (Fig. 3F). Additional experiments showed that the FRYL C-terminus (aa 2605-3013) was sufficient for Homer binding and that mutations in the first (PSPF) or the second (PTVF) Homer binding site completely disrupted the interaction, indicative of a cooperative binding mechanism (Fig. S5E and S5F). Furthermore, Homers did not co-IP with Furry, confirming the specificity of the interaction. We concluded that FRYL interacts directly and specifically with Homers via two PxxF motifs.

To explore a potential role of FRYL in YAP signaling, we produced *FRYL* MDCK KO cells. Interestingly, loss of FRYL reduced YAP S127 phosphorylation and increased both TEAD reporter activity and YAP nuclear translocation in at least one out of the two KO clones examined (Fig. 3G-3I). Moreover, genetic inactivation of *FRYL* in MDCK cells as well as siRNA-mediated silencing of FRYL in 293T or HCT116 cells significantly increased YAP target gene transcription (Fig. 3J and Fig. S5G), demonstrating that FRYL acts as a robust suppressor of YAP/TEAD signaling. By contrast, the expression of the Wnt-associated genes FZD4, GNA14, DACH1 and HAS2 was reduced in *FRYL* KO cells (Fig. 3K). Axin2 mRNA levels were also decreased but Wnt reporter activity was unexpectedly unchanged (Fig. 3L and 3M). This indicates that Homers and FRYL function antagonistically in regulating the transcription of canonical YAP/TEAD target genes but synergize to control Wnt target genes and pathway regulators.

### Homers form biomolecular condensates in 293T and HCT116 cells

Previous studies demonstrated that Homers act as scaffolds for the assembly of biomolecular condensates (47, 48). Biomolecular condensates are membrane-less compartments that concentrate macromolecules via multivalent interactions, generating phase-separated liquid-like or gel-like assemblies that vary in how rapidly they exchange materials with the surrounding environment (91, 92). To determine whether Homer phase separation contributes to YAP and Wnt signaling, we analysed the localization and dynamics of Homers in 293T cells. Interestingly, transient overexpression of GFP-tagged Homers in 293T cells produced large cytoplasmic puncta (Fig. 4A and S6A). Live cell imaging revealed that these structures emerge from small nucleation sites and grow through fusion and coalescence (Fig. S6C and S6D). The puncta were devoid of a surrounding membrane and displayed considerable ultrastructural heterogeneity, as determined by transmission electron microscopy (Fig. S6E and S6F). In addition, Fluorescence Recovery After Photobleaching (FRAP) analysis showed that Homers are largely immobile within these assemblies, with only 13-26% of Homer fluorescence recovery within 1 min after photobleaching (Fig. S6G). This indicated that Homers form condensates when overexpressed in cells.

**Figure 4:**
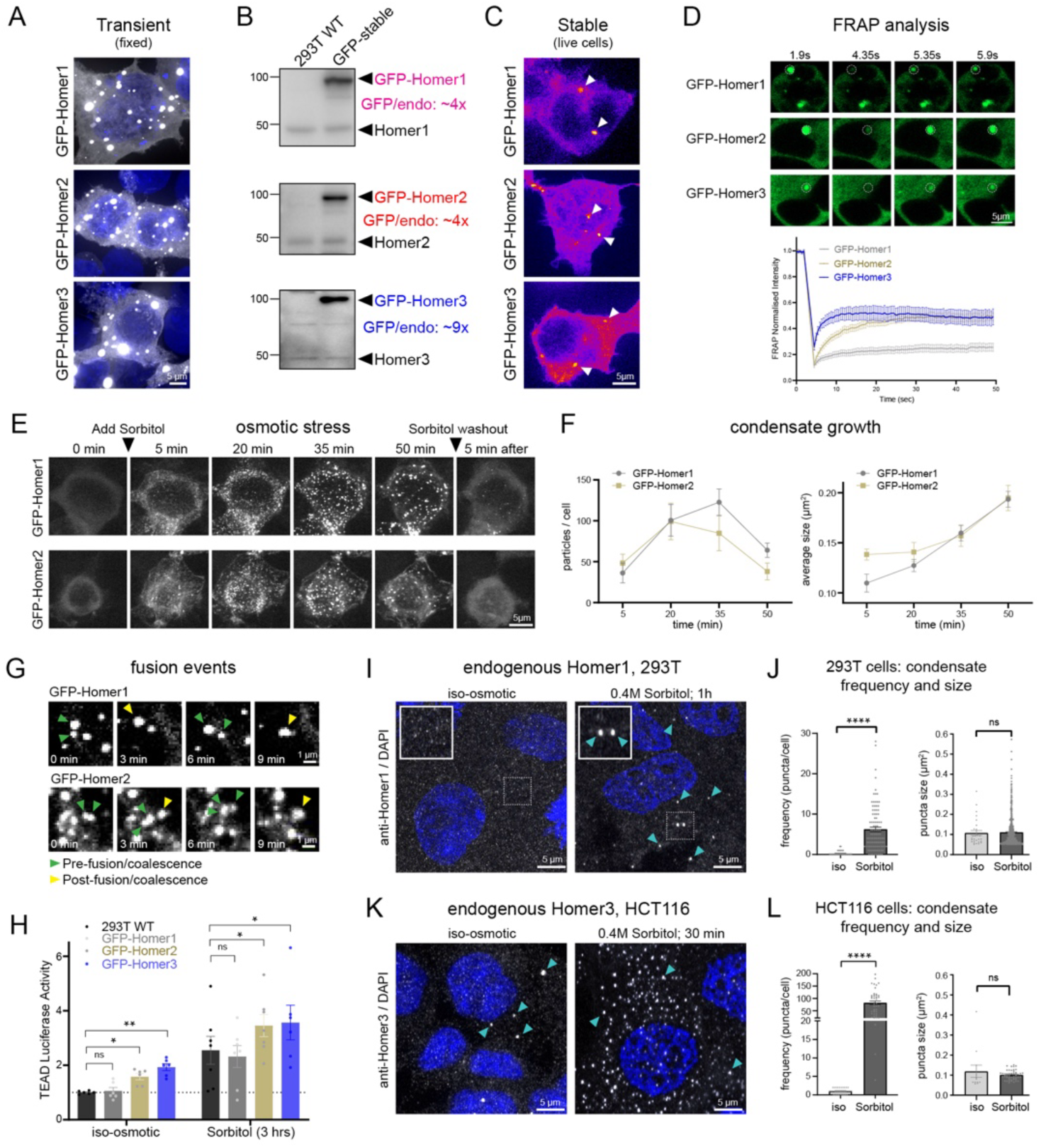
Homers form cytoplasmic biomolecular condensates at endogenous levels. (A) Confocal micrographs of fixed 293T cells transiently transfected with GFP-Homer1, 2, or 3. (B) Western blot analysis of 293T cells stably transfected with GFP-Homer1, 2, or 3. The degree of overexpression was determined by densitometry as GFP-Homer over endogenous Homer signal (n=3). (C) 293T cells stably transfected with GFP-Homer1, 2, and 3 imaged live by spinning disk microscopy. Note the presence of small cytoplasmic puncta. (D) FRAP analysis of GFP-Homer1, 2, and 3 puncta in stably transfected 293T cells (n=20). (E) Stable GFP-Homer cells were imaged live by spinning disk microscopy before and after the addition of 0.4 M sorbitol for 1 hour. (F) Quantification of particle size (area) and frequency before and after induction of hyperosmotic shock based on data shown in (E) (n=10). (G) Fusion events of GFP-Homer1 and GFP-Homer2 puncta. (H) TEAD luciferase assay in wild-type 293T cells and stable GFP-Homer cell lines grown in isosmotic medium or in medium containing 0.4 M sorbitol for 3 hours. Note that overexpression of GFP-Homer2 or GFP-Homer3 enhances TEAD activity under isosmotic and hyperosmotic conditions. (I) Airyscan microscopy of fixed 293T cells grown in isosmotic medium or in medium containing 0.4 M sorbitol for 1 hour stained with Homer1 antibodies. (J) Quantification of Homer1 puncta size and frequency based on data shown in (I). (K) Airyscan microscopy of fixed HCT116 cells grown in isosmotic medium or in medium containing 0.4 M sorbitol for 30 min stained with Homer3 antibodies. (L) Quantification of Homer3 puncta size and frequency based on data shown in (K). Data in J and L is presented as mean ± SEM. ****P ≤ 0.0001 (Student’s t-test; n=33-50 cells each).

To determine if Homers phase separate at or close to physiological expression levels, we generated stable 293T cell lines expressing GFP-Homer1 and GFP-Homer2 at ∼4 fold above endogenous levels and GFP-Homer3 at ∼9 fold over baseline (Fig. 4B). All GFP-Homer proteins produced cytoplasmic puncta in live cells, though at markedly lower frequency than after transient overexpression (Fig. 4C). FRAP analysis revealed substantially greater mobility for GFP-Homer2 and GFP-Homer3 (∼50% recovery within 20 sec upon photobleaching), whereas GFP-Homer1 remained largely immobile (Fig. 4D). This suggests that Homers overexpressed at high levels assemble condensates with gel-like characteristic, whereas those formed at close to endogenous expression levels tend to exhibit more liquid-like properties.

Hyperosmotic stress (or macromolecular crowding) has been shown to promote the phase separation of Hippo signaling molecules (11, 14) and enhance YAP/TEAD activity (12, 93). Interestingly, addition of 0.4 M sorbitol to live 293T cells stably expressing GFP-Homers triggered a rapid and fully reversible increase in the number of cytoplasmic puncta (Fig. 4E, MOVIE S1). Over time, the number of puncta decreased while their average size increased, consistent with coarsening (Fig. 4F). Fusion events between individual puncta were frequently observed (Fig. 4G, MOVIE S2), supporting their liquid-like and dynamic behavior.

To test whether Homer condensates promote YAP/TEAD signaling, we performed TEAD luciferase assays in wild-type and stable GFP-Homer 293T cells grown under isosmotic or hyperosmotic conditions. Osmotic stress enhanced TEAD activity in wild-type 293T cells, as expected (12, 93) (Fig. 4H). In line with our data from transient overexpression (Fig. S4D), stable overexpression of GFP-Homer2 or GFP-Homer3 (but not GFP-Homer1) significantly increased TEAD activity under basal conditions, and a further increase was observed in response in sorbitol-treated cells (Fig. 4H). We concluded that Homer proteins promote YAP/TEAD signaling through phase separation.

Next, we examined endogenous Homer1 in fixed 293T cells using Airyscan microscopy (which offers a lateral resolution of ∼100-120 nm). Under isosmotic conditions, Homer1 immunostaining appeared predominantly diffuse and cytoplasmic puncta were rarely detected. Intriguingly, exposure of cells to osmotic stress markedly increased the number of Homer1 puncta (Fig. 4I). These structures were ∼350 nm in diameter (average area ∼0.1 µm²) (Fig. 4J) and therefore similar in size to the puncta observed in sorbitol-treated GFP-Homer–expressing cells (diameter ∼500 nm; average area ∼0.2 µm²) (Fig. 4F). Furthermore, immunostaining of Homer3 in HCT116 cells (which express substantially higher levels of Homer3 than 293T cells (Fig. S4G)) revealed distinct cytoplasmic puncta even under isosmotic conditions (Fig. 4K), and the frequency of these puncta increased dramatically in response to osmotic stress (Fig. 4L). Taken together, these data demonstrate that Homer proteins phase separate at endogenous expression levels.

### FRYL and PATJ differentially regulate Homer phase separation

Given that YAP can phase separate with Hippo pathway proteins in the nucleus and the cytoplasm (7–14), we wondered whether Homer condensates sequestered YAP. We found that YAP-GFP colocalized with Homers when both proteins were co-overexpressed (Fig. S7A and S7B). However, endogenous YAP was not detected in Homer condensates induced by osmotic stress (Fig. S7C and S7D). In addition, YAP-GFP condensates induced by osmotic stress did not colocalize with endogenous Homers (Fig. S7E). This indicates that, under physiological conditions, YAP is largely excluded from Homer condensates, and that Homer-mediated YAP activation does not involve YAP sequestration.

Next, we addressed the functions of FRYL and PATJ in Homer phase separation. Interestingly, transient overexpression of GFP-FRYL, but not the FRYL HBM, resulted in the spontaneous formation of cytoplasmic droplet-like condensates (Fig. S6B). Endogenous Homer1 was efficiently recruited into these structures (Fig. 5A), indicating that FRYL promotes Homer phase separation even in the absence of osmotic stress. GFP-FRYL also colocalised with mCherry-Homer3 in live cells (Fig. 5B and MOVIE S3) and with HA-tagged Homer1 in fixed cells (Fig. 5C), and colocalization was significantly reduced when the two Homer binding sites in FRYL were mutated (Fig. 5D and 5E). We further found that Homer1 condensates were significantly larger in cells co-transfected with wild-type FRYL compared to cells co-transfected with the FRYL HBM (Fig. 5F). Moreover, FRAP analysis demonstrated that wild-type (but not mutant) FRYL significantly increased the mobile fraction of Homer3 from ∼25% to ∼36% (Fig. 5G). FRYL itself exhibited minimal fluorescence recovery (∼10% mobile fraction), indicating a tight association with the Homer scaffold. We concluded that FRYL promotes the phase separation of endogenous Homers in a PxxF motif-dependent manner.

**Figure 5:**
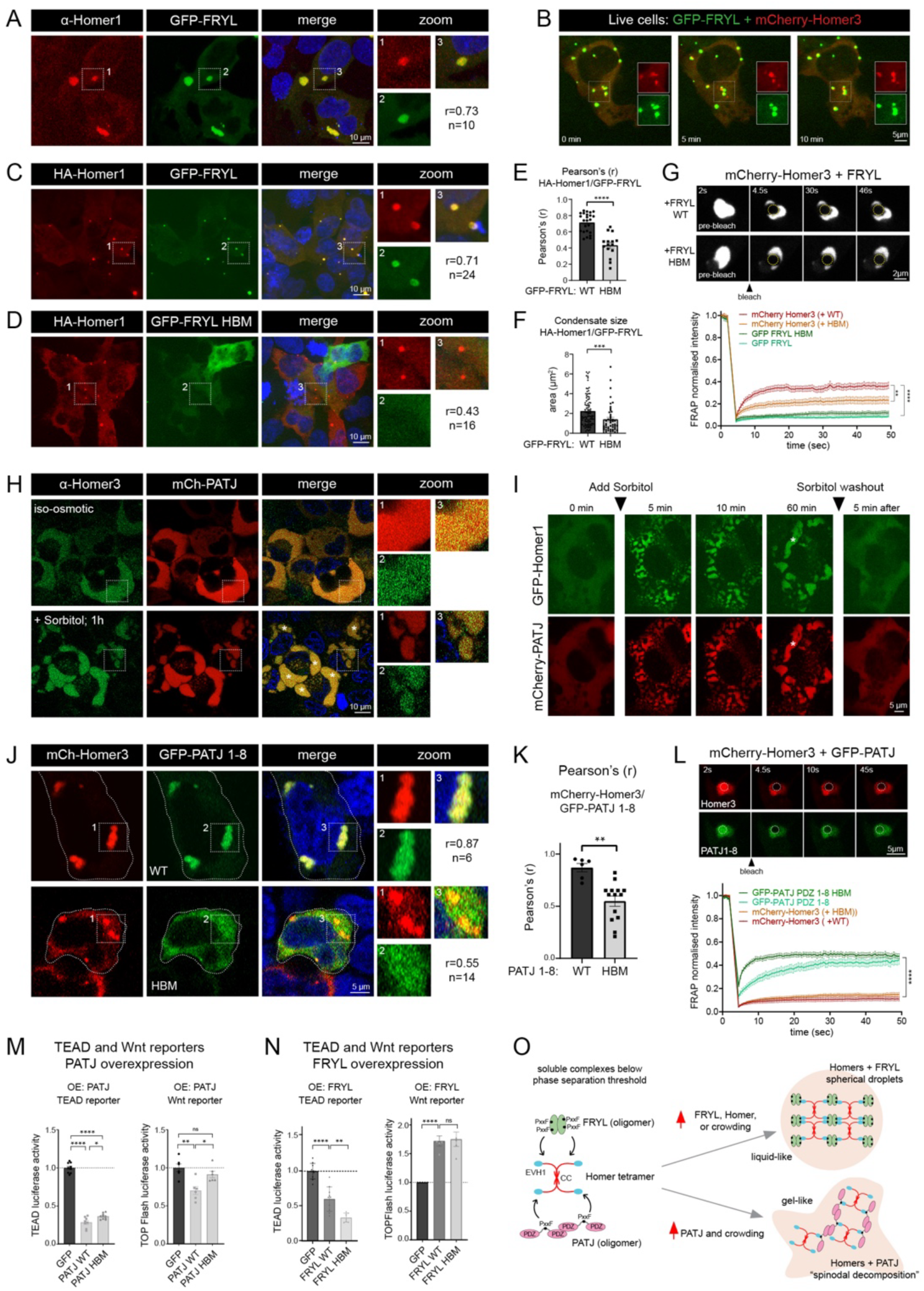
FRYL and PATJ differentially regulate Homer phase separation. (A) 293T cells transiently transfected with GFP-FRYL were fixed and stained with Homer1 antibodies. (B) Live cell microscopy of 293T cells co-transfected with mCherry-Homer3 and GFP-FRYL. (C and D) 293T cells were co-transfected with HA-Homer1 and GFP-FRYL (C) or the GFP-FRYL HBM M2 (D). (E) Pearson’s correlation (r) analysis of HA-Homer1 and GFP-FRYL vs. HA-Homer1 and GFP-FRYL HBM M2. (F) Size measurements of HA-Homer1 condensates in cells co-transfected with GFP-FRYL or GFP-FRYL HBM M2. (G) FRAP analysis of mCherry-Homer3 in 293T cells co-transfected with either GFP-FRYL or GFP-FRYL HBM M2 (n=17). (H) 293T cells transiently transfected with mCherry-PATJ grown under isosmotic or hyperosmotic conditions for 1 hour were fixed and stained with Homer3 antibodies. (I) Live cell microscopy of stable GFP-Homer1 293T cells transfected with mCherry-PATJ before and after sorbitol addition, and after sorbitol washout. (J) 293T cells transiently co-transfected with mCherry-Homer3 and either GFP-PATJ PDZ1-8 or GFP-PATJ PDZ1-8 HBM. (K) Pearson’s r co-localization analysis of mCherry-Homer3 with WT and mutant GFP-PATJ PDZ1-8. Data is presented as mean ± SEM. Statistical significance was assessed using Student’s t-test (n=15). (L) FRAP analysis of 293T cells co-transfected with mCherry-Homer3 and either GFP-PATJ PDZ1-8 or the corresponding HBM (n=10). (M) TEAD and TOPFlash reporter assays of 293T cells transfected with GFP, WT GFP-PATJ PDZ1-8, or GFP-PATJ PDZ1-8 HBM (n=4). (N) TEAD and TOPFlash reporter assays of 293T cells transfected with GFP, WT GFP-FRYL, or GFP-FRYL HBM M2 (n=5). FRAP data in (G) and (L) is presented as mean ± SEM. Statistical analysis was performed using two-way ANOVA. Luciferase assay data were analysed using a paired Student’s t-test. * P ≤ 0.05, ** P ≤ 0.01, *** P ≤ 0.001, **** P ≤ 0.0001. (O) Model of how osmotic stress and changes in Homer, PATJ, and FRYL expression modulate Homer phase separation (see discussion). FRYL is depicted as a hypothetical dimer but could equally enhance valency as a monomer (due to its two Homer binding sites). PATJ is depicted as PDZ5-6 fragment for simplicity and may oligomerise via PDZ-PDZ domain interactions.

In contrast to FRYL, overexpressed mCherry-PATJ did not induce the formation of Homer condensates under isosmotic conditions (Fig. 5H, top panel). Interestingly, however, upon osmotic stress, mCherry-PATJ and stably expressed GFP-Homer1 rapidly coalesced into irregularly shaped networks, which progressively merged into large, micrometer-size cytoplasmic patches (Fig. 5I, MOVIE S4). These structures contained endogenous Homers (Fig. 5H, bottom panel) and dissolved rapidly upon sorbitol removal (Fig. 5I), indicating a dynamic and reversible condensation mechanism. We then co-expressed PATJ PDZ1-8 with mCherry-Homer3 and found that this resulted in the appearance of more spherical, droplet-like condensates (Fig. 5J, MOVIE S5). As expected, mutation of the Homer-binding site in PATJ PDZ1-8 markedly impaired its incorporation into these condensates and significantly reduced the colocalization with Homer3 (Fig. 5J and 5K). FRAP analysis further revealed that PATJ was relatively mobile and did not alter Homer3 mobility (Fig. 5L), distinguishing its behavior from that of FRYL. These findings indicate that FRYL and PATJ differentially modulate Homer phase separation, giving rise to condensates with distinct morphologies and material properties.

These findings prompted us to examine whether the effects of PATJ and FRYL on YAP/TEAD and Wnt signaling depend on their direct interaction with Homers. Interestingly, overexpression of PATJ PDZ1-8 in 293T cells significantly reduced both TEAD and TOPFlash activity, and mutation of the Homer binding site partially (TEAD) or completely (TOPFlash) reversed this effect (Fig. 5M). Overexpression of FRYL also reduced TEAD reporter activity, but its inhibitory effect was significantly enhanced when the Homer binding site in FRYL was inactivated (Fig. 5N). TOPFlash reporter activity was increased upon overexpression of FRYL, independently of its association with Homers (Fig. 5N). Collectively, the data demonstrate that PATJ and FRYL modulate Homer-dependent signaling outputs through direct PxxF motif–mediated interactions, while also revealing interaction-dependent and -independent effects on YAP and Wnt pathway activity.

## Discussion

Biomolecular condensates have emerged as key regulators of signal transduction, including the Hippo/YAP and Wnt/β-catenin pathways (8, 92, 94–96). Here, we identify Homer proteins as polarity-associated scaffolds that phase separate in epithelial cells and coordinate YAP and canonical Wnt signaling. We further show that the material properties and signaling outputs of Homer condensates are modulated by the Crumbs complex component PATJ and the NDR scaffold FRYL. Together, our data support a model in which Homer condensates function as a tunable regulatory hub linking epithelial polarity cues to YAP–Wnt pathway crosstalk.

### Homer phase separation as a signaling control mechanism

Homers are multivalent scaffolds that oligomerize via coiled-coil domains and bind EVH1 domain ligands containing PxxF motifs (45, 46, 97). In neurons, these interactions drive phase separation with postsynaptic density proteins to regulate actin organization and synaptic signaling (47–49). Our findings extend this principle to epithelial cells, where Homers form condensates at or near endogenous expression levels. These assemblies exhibit hallmark features of phase-separated compartments, including rapid molecular exchange, fusion and coarsening-driven growth, and reversible assembly under hyperosmotic stress. Thus, Homers appear to operate near their phase boundary, allowing modest changes in expression, ligand availability, or macromolecular crowding to trigger condensation (Fig. 5O).

Despite high sequence similarity, Homer1, Homer2, and Homer3 formed condensates with distinct ultrastructural and material properties. Their effects on YAP/TEAD signaling were similarly heterogeneous and cell-type dependent. This indicates that relative abundance and subtle differences in ligand binding or oligomerization dynamics can alter Homer network connectivity, reshaping both condensate material state and signaling output (98, 99). Indeed, strong Homer overexpression promoted more gel-like assemblies capable of sequestering YAP, whereas fluid condensates formed at near-endogenous levels did not. Homer dosage therefore correlates with condensate material properties and YAP recruitment. This dosage sensitivity may be particularly relevant in disease. Homers—especially Homer3—are upregulated in multiple tumor types and correlate with aggressive behavior and poor prognosis (54–56). Elevated Homer expression could shift condensates toward more stable or aberrantly interconnected states, altering YAP activity and contributing to tumor progression (8).

### PATJ and FRYL differentially tune the Homer condensate network and YAP signaling

We find that PATJ and FRYL both associate with endogenous Homers in a PxxF motif-dependent manner but exert strikingly distinct effects on condensate assembly and dynamics (Fig. 5O). FRYL, which contains two PxxF motifs, acted as a multivalent co-scaffold: its overexpression drove droplet formation even in the absence of osmotic stress and increased Homer mobility despite being itself relatively immobile. This suggests that FRYL enhances network connectivity while simultaneously fluidizing Homer condensates. In contrast to FRYL, PATJ did not nucleate condensates under isosmotic conditions. Intriguingly, however, upon osmotic stress, PATJ rapidly demixed with Homers into irregular, interconnected cytoplasmic networks that coarsened over time and fully dissolved upon sorbitol washout. The absence of discrete nucleation events and the emergence of bi-continuous, non-spherical domains are consistent with a spinodal decomposition-like phase separation mechanism (100). Moreover, when co-expressed with Homers, PATJ remained relatively mobile and did not significantly alter Homer mobility. Taken together this suggests that PATJ acts as a multivalent but dynamic crosslinker that increases network connectivity, thereby shifting the material state toward a percolated, gel-like phase. Because PATJ contains a single Homer-binding motif, cooperative network assembly likely depends on PATJ oligomerization via PDZ-PDZ interactions or associated binding partners (101) (Fig. 5O). Although we focused on PATJ and FRYL, we note that additional scaffolds and clients are expected to influence Homer condensate composition and material properties (100, 102, 103). Given that Hippo pathway regulators modulate YAP through cytoplasmic phase separation (9, 11, 14), it will be important to determine whether Homer condensates are related to these assemblies or whether they constitute a distinct type of biomolecular condensate.

Functionally, we show that PATJ, Homers, and FRYL regulate YAP/TEAD (and Wnt, see below) signaling through direct inhibitory interactions. We propose that Homer condensates sequester and suppress the FRYL/NDR complex, thereby limiting NDR-mediated YAP phosphorylation and promoting YAP/TEAD-dependent transcription. Although a direct role of NDR downstream of Homers and FRYL has yet to be established, this interpretation is consistent with reports showing that NDR phosphorylates and inhibits YAP in the mouse intestine and in cultured cells (4, 5). PATJ, in turn, appears to counteract Homer function and thereby limit YAP-driven transcription. However, PATJ’s role is likely complex as its inactivation in MDCK cells enhanced endogenous YAP target gene expression while simultaneously reducing TEAD reporter activity. Although the precise signaling mechanisms have yet to be defined, PATJ might modulate YAP target gene transcription through TAZ, through alternative transcription factors, such as AP-1 or SMAD2/3 (40, 104–106), or through additional binding partners, such as AMOT and KIBRA (58, 60, 107).

Collectively, our findings suggest that PATJ acts as a polarity-dependent brake on Homer-driven YAP activation. This agrees with the well-established role of the Crumbs complex in linking cell density and polarity cues to the inhibition of YAP (32, 39, 40). As a core Crumbs complex component, PATJ may recruit Homers to the apical-lateral border, thereby restricting their ability to activate YAP. Given that PATJ promotes ZO-1 condensation at tight junctions (61) and that membrane surfaces can facilitate phase separation (108, 109), it is plausible that PATJ dampens Homer-mediated YAP activation by modulating Homer condensation at apical cell junctions.

### Homer condensates coordinate YAP-Wnt crosstalk

Our data further indicate that the PATJ–Homer–FRYL axis links YAP regulation to canonical Wnt/β-catenin signaling. Importantly, downregulation of Homers in HCT116 cells suppressed both YAP/TEAD and Wnt pathway activity and reduced cell migration. This is consistent with a reciprocal positive feedback loop between YAP and Wnt in colorectal cancer cells (22, 23, 25, 82) and suggests a tumor-promoting role for Homers in Wnt-driven cancers.

Interestingly, while Homers and FRYL acted antagonistically on classical YAP target genes, both enhanced Wnt reporter activity and the expression of several pro-oncogenic, Wnt-associated genes, including FZD4, GNA14, DACH1, and HAS2 (70–73, 78, 79, 81, 110–112). Whilst FRYL-mediated suppression of YAP was modulated by its interaction with Homers, its Wnt-promoting activity was not. This suggests that Homers and FRYL promote a Wnt-related transcriptional program independently of their opposing roles in YAP signaling. We propose, therefore, that Homers and FRYL converge to fine-tune YAP activity but influence β-catenin output via distinct, YAP-independent mechanisms (Fig. 6). In non-polarised or cancer cells, Homer assemblies may sequester and suppress the FRYL/NDR complex, thereby promoting YAP activation. In parallel, Homers may enhance β-catenin signaling through a Src-dependent pathway (56). Loss or polarity-dependent inactivation of Homers increases YAP phosphorylation and cytoplasmic retention, dampening Wnt signaling via YAP-β-catenin crosstalk (20, 21, 23) and by eliminating direct Src-mediated β-catenin activation. Conversely, FRYL depletion enhances YAP/TEAD transcription by reducing NDR activity but diminishes Wnt pathway output. Because NDR has been reported to promote β-catenin degradation (113, 114), FRYL’s pro-Wnt effect is unlikely to be mediated by its NDR scaffolding function. Altogether our and previous work suggest that NDR may operate in functionally distinct pools: a FRYL-associated pool that regulates YAP activity, and a separate pool that controls β-catenin turnover and Wnt signaling independently of FRYL.

**Figure 6:**
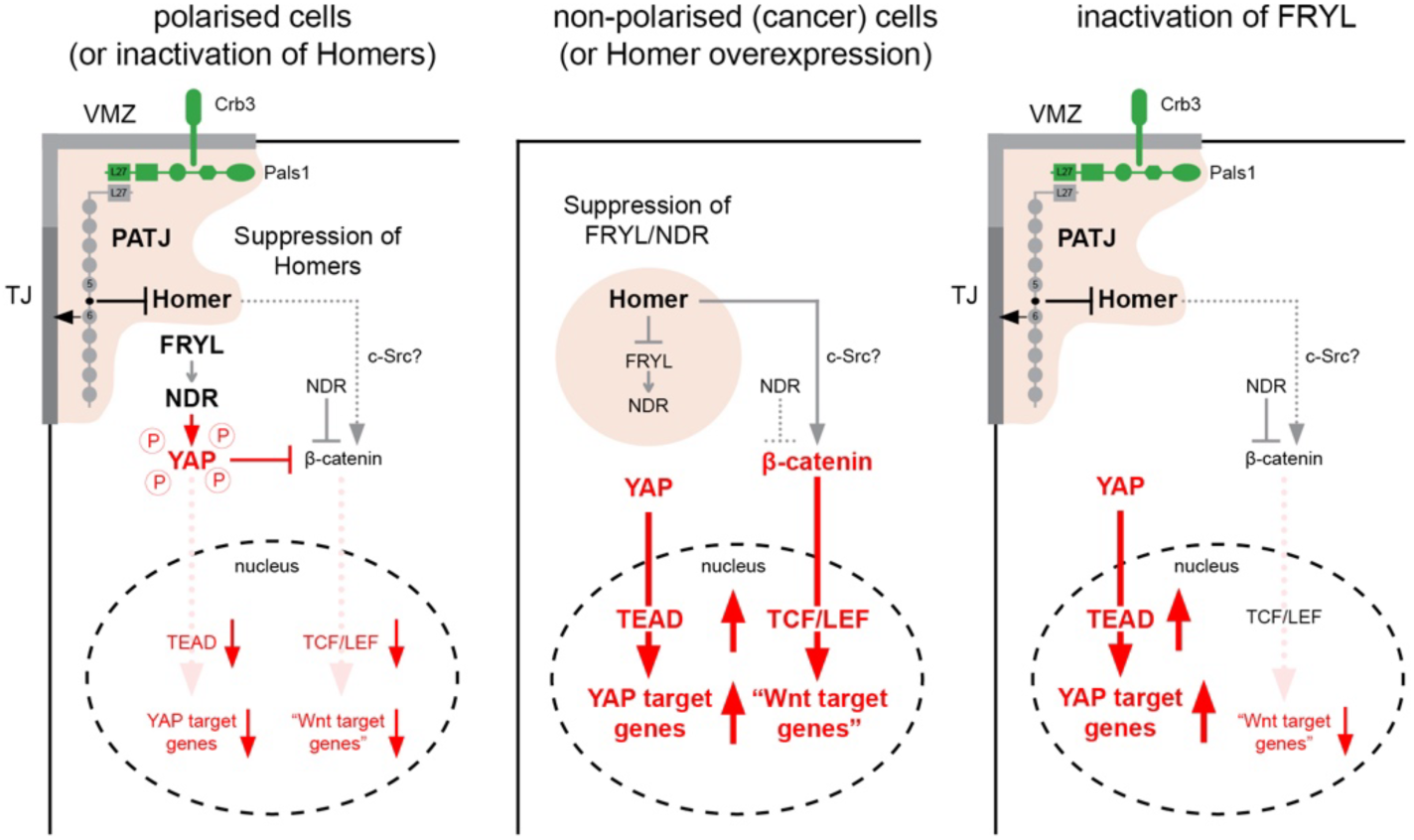
Model of Homer-mediated YAP-Wnt signaling crosstalk. Hypothetical mechanism of how PATJ, Homers, and FRYL control YAP-Wnt signaling crosstalk (see discussion). The model assumes inhibitory interactions between PATJ and Homers and between Homers and the FRYL/NDR complex. FRYL suppresses YAP via NDR, and NDR functions independently of FRYL to suppress β-catenin.

Collectively, our findings support a model in which Homer phase separation constitutes a tunable signaling platform integrating polarity cues with Hippo and Wnt pathway activity. In polarized epithelial cells, PATJ and the Crumbs complex restrain Homer-driven YAP activation by recruiting it to the apical-lateral border. In cancer cells—where Homers are often upregulated and polarity is disrupted—this restraint may be lost. Elevated Homer expression and altered condensate properties may shift signaling toward persistent YAP and Wnt activation, promoting migration and tumor progression. Homer condensates therefore function as dynamic regulatory hubs that may represent targets for modulating oncogenic YAP and β-catenin signaling in epithelial-derived cancers.

## Materials & Methods

### Cell lines

Madin-Darby Canine Kidney type II (MDCK-II), Human Embryonic Kidney 293T (HEK293T), and three human colorectal carcinoma cell lines (HCT116, DLD-1, and RKO) were used in this study. MDCK-II, HEK293T and HCT116 cells were grown in DMEM, high glucose, GlutaMAX™ pyruvate (Gibco #10569010). DLD-1 and RKO cells were grown in RPMI 1640 supplemented with L-Glutamine (Gibco #11875093). All media were supplemented with 10 % foetal bovine serum and 100 U/ml penicillin-streptomycin. All cells were grown at 37 ^O^C and under 5% CO_2_. Cell lines used were not authenticated and were negative to mycoplasma contamination. For some experiments MDCK cells were seeded onto polycarbonate or polyethylene 0.4 µm-pore Transwell® filter inserts at a seeding density of approximately 2.2×10^5^ cells/cm^2^. Cells were grown on the filter for 10-14 days to achieve full polarisation. Media was replenished every two days.

### Bacterial strains

NEB10-beta competent *E. coli* (NEB, #C3019I) were used to transform plasmids for cloning. Bacteria were cultured in LB Agar (BD, Difco #244520) and LB Broth (BD, Difco #244620) at 37°C.

### DNA constructs

Homers, PATJ, FRYL, and NDR1 cDNAs were PCR amplified from commercial vectors, utilizing primers specifically designed to enable seamless ligation into the pEGFP-C1, p3xHA-C1, or pEGFP-APEX2-C1 backbones (43). PATJ and NDR1 constructs were generated using Transfer PCR (TPCR), using www.rf-cloning.org/ for primer design. Details of cDNA cloning and primers are listed in Table S1. Full-length FRYL was cloned into the pEGFP-C1 backbone using Gibson Assembly HiFi DNA Assembly Cloning Kit (NEB #E5520). Site-directed mutagenesis PCR was used to introduce specific mutations into the wild-type cDNAs. For PATJ, mutations P812A and F815A were introduced to inactivate the PxxF motif in the linker region between PDZ domains 5 and 6. Homer1 was mutated at position G89A, rendering the EVH1 domain inactive. The two PxxF motifs in FRYL were mutated to produce the mutants M1 (P2663A & P2757A) and M2 (P2663A & F2666A / P2757A & F2760A). The specific cDNAs and primers used for these mutagenesis reactions are detailed in Table S1. CRISPOR (http://crispor.org) was used to design gRNA. Three gRNAs targeting early exons were selected for each gene of interest. The gRNAs were synthesized as pairs of short oligonucleotides with 5’ phosphorylated ends. These oligonucleotide pairs were annealed in annealing buffer (1 mM Tris pH 8, 5 mM NaCl, 0.5 mM EDTA in Milli-Q water) by heating the mixture to 95°C, followed by a gradual cooling to 30°C for 1 hour. The annealed gRNAs were then ligated into pSpCas9(BB)-2A-Puro (PX459) (Addgene #62988) vector digested with *BbsI*-HF® (NEB). All gRNAs used in this study are listed in Key Resource Table. All plasmids were transformed into NEB10β (NEB #C3019I) competent *E.coli*. All plasmids were screened by restriction enzyme digest or PCR and verified by DNA sequencing.

### Plasmid and gRNA transfection

MDCK and HEK293T cells were transfected with plasmids encoding GFP- or HA-tagged proteins, and/or guide RNAs (gRNAs), when cells reached approximately 60% confluency. Prior to transfection, the culture medium was replaced with fresh antibiotic-free DMEM containing 10% FBS. Transfections were carried out using either polyethylenimine (PEI) or Lipofectamine™ 3000 (Invitrogen #L3000015), depending on the cell type. For MDCK cells, Lipofectamine 3000 was used at a ratio of 3 µL per 1 µg of plasmid DNA. For HEK293T cells, PEI was used at a ratio of 5 µL per 1 µg of plasmid DNA. In 6-well plates, a total of 2 µg DNA was used per well. For transfection in 12-well or 24-well plates, 1 µg DNA per well was used for MDCK cells. DNA and transfection reagents were diluted in DMEM or OptiMEM and incubated at room temperature for 30 minutes to allow complex formation. The transfection mix was then added dropwise to the cells. After 4 hours, the medium was replaced with complete growth medium supplemented with 10% FBS and 1% penicillin–streptomycin. Cells were allowed to grow for 16-24 hours post-transfection before being processed for downstream RNA or protein analyses.

### Generation of MDCK knockout cells

Stable MDCK cell lines expressing EGFP-APEX2 tagged Homers (43) and PATJ knockout MDCK cells (64) have been described previously. To generate Homer and FRYL knockout cells, MDCK cells were co-transfected with 0.5 µg of PX459 plasmid (encoding Cas9 and the single-guide RNA targeting the gene of interest) and 0.1 µg of pEGFP-C1. The pEGFP-C1 plasmid encoded a neomycin (G418/geneticin) resistance cassette, allowing for antibiotic selection. 24 hours after transfection, cells were placed under 200 µg/mL G418 selection for 14 days. Surviving cells were then sorted by FACS based on GFP expression and plated as single-cell clones in 96-well plates. Clonal populations were expanded and screened for knockout efficiency by Western blotting to confirm the absence of the target protein. Knockout clones were further validated by Sanger sequencing of the targeted locus and by immunofluorescence staining to confirm the loss of protein expression at the cellular level.

### Genomic DNA Extraction and Sanger Sequencing

Genomic DNA (gDNA) was isolated from cultured cells using the GeneJET Genomic DNA Purification Kit (Thermo Scientific #K0721) according to the manufacturer’s instructions. Briefly, cells were harvested and washed twice with PBS. The resulting cell pellet was lysed in a buffer containing proteinase K to degrade cellular proteins, followed by RNase A treatment to eliminate RNA contamination. After complete lysis, ethanol was added to facilitate DNA binding to the silica membrane within the spin column. The lysate was then transferred to the spin column and centrifuged, allowing gDNA to bind to the membrane while contaminants were removed via a series of washes with the supplied buffers. The purified DNA was eluted in RNase-free water. For genotyping of CRISPR-edited loci, PCR amplification was performed using primers flanking the guide RNA target site. Primer sequences were designed using the CRISPOR online tool (http://crispor.tefor.net) to ensure specificity. PCR products were purified and submitted for Sanger sequencing (BioBasic, Singapore). Resulting chromatograms were analyzed to confirm sequence edits and to assess the efficiency of gene knockout.

### siRNA transfection

For siRNA transfection, cells were seeded to reach approximately 50% confluency on the day of transfection. Culture medium was replaced with antibiotic-free DMEM containing 10% FBS. siRNA duplexes were transfected at a final concentration of 100 nM using Lipofectamine RNAiMAX (Invitrogen, #13778075). For every 100 nM of siRNA, 3 µL of RNAiMAX was used per well. siRNA and RNAiMAX were diluted in DMEM only media, incubated for 20–30 minutes at room temperature, and then added dropwise to cells. After 24 hours, the medium was replaced with fresh complete DMEM containing FBS and antibiotics. Cells were harvested for RNA or protein extraction 48 hours post-transfection.

### Immunoprecipitation

MDCK or HEK293T cells transiently transfected with GFP-tagged constructs were washed twice with cold phosphate-buffered saline (PBS) and lysed in ice-cold lysis buffer composed of 50 mM Tris-HCl (pH 7.4), 150 mM NaCl, 0.5% Triton X-100, and 1 mM EDTA, supplemented with protease inhibitors (Roche cOmplete™ Mini EDTA-free, #11873580001), and, where indicated, phosphatase inhibitors (PhosSTOP™, Roche). Lysis was performed by incubating the cells with buffer for 10 minutes at 4 °C on a rotating platform. Lysates were clarified by centrifugation at 20,000 × g for 30 minutes at 4 °C. Cleared lysates were incubated with GFP-Trap® Agarose beads (Chromotek, gta-20) at a 1:20 bead-to-lysate dilution in lysis buffer. The bead–lysate mixture was incubated at 4 °C for 2 hours on a rotator to allow binding of GFP-tagged proteins. Following incubation, beads were pelleted by centrifugation at 500 × g for 2 minutes at 4 °C and washed three times with lysis buffer to remove unbound proteins. After the final wash, beads were resuspended in 4× LDS sample buffer (Invitrogen) containing 200 mM dithiothreitol (DTT). Samples were boiled at 96 °C for 5 minutes to elute bound proteins from the beads.

### SDS-PAGE and Western Blotting

Protein lysates were mixed with 4× lithium dodecyl sulfate (LDS) sample buffer containing 200 mM dithiothreitol (DTT) and denatured by boiling at 96 °C for 5 minutes. Total protein concentrations were determined using the Bradford assay (Bio-Rad) to ensure equal loading across samples. Equal amounts of protein were separated by SDS-PAGE at 120 V for 1 hour in standard SDS running buffer (25 mM Tris base, 192 mM glycine, 0.2% SDS). Proteins were transferred onto methanol-activated polyvinylidene difluoride (PVDF) membranes (Millipore) in transfer buffer (25 mM Tris, 192 mM glycine, 20% (v/v) methanol) using a wet-transfer system, either overnight at 20 V or for 3 hours at 70 V, both at 4 °C. After transfer, membranes were briefly dehydrated in methanol for 10 seconds and air-dried for 30 minutes to enhance protein retention. Membranes were then blocked and incubated with primary antibodies diluted in blocking buffer containing 2.5% BSA, 0.4% Tween-20, and 1 mM sodium azide. Following three washes with PBS containing 0.1% Tween-20 (PBST), membranes were incubated with species-appropriate horseradish peroxidase (HRP)-conjugated secondary antibodies diluted in 5% non-fat dry milk in PBST. After a further three washes in PBST, blots were developed using an enhanced chemiluminescent (ECL) substrate and imaged using a ChemiDoc imaging system (Bio-Rad).

### Immunofluorescence

HEK293T cells were seeded onto fibronectin-coated glass coverslips to promote robust adhesion. MDCK cells were cultured either on glass coverslips or on Transwell® filters, depending on the experimental setup. Cells were washed three times with PBS++ (PBS supplemented with Ca²⁺ and Mg²⁺) and fixed with either 1% or 4% paraformaldehyde (PFA), depending on primary antibody used, in PBS++ for 10-20 minutes at room temperature, followed by three additional washes with PBS++. Cells were then permeabilised with 0.5% Triton X-100 in PBS for 10-15 minutes, washed three times in PBS, and subsequently blocked in 10% FBS in PBS for 1 hour at room temperature. For Transwell-grown monolayers, the membrane filters were carefully excised using sterile razor blades and handled with the cell-facing side up throughout all staining steps. All incubation steps, including washes, were performed in a humidified chamber to prevent drying.

Samples were incubated with primary antibodies diluted in antibody incubation buffer (0.2 µm-filtered PBS containing 0.1% BSA and 0.01% Tween-20) for 1–3 hours at room temperature. After three PBS washes, samples were incubated with fluorophore-conjugated secondary antibodies diluted in the same buffer for 1 hour at room temperature, protected from light. Following secondary antibody incubation, samples were stained with 20 ng/mL DAPI in PBS to label nuclei and washed thoroughly. Glass coverslip samples were mounted onto microscope slides using fluorescent mounting medium (Vectashield, Vector Laboratories H-1000). For Transwell® samples, the membranes were placed cell side up, overlaid with mounting medium, and covered with #1.5 (12 mm) glass coverslips (VWR). Coverslips were sealed using transparent nail polish to prevent drying and preserve fluorescence.

### Confocal Microscopy

Fixed samples and live-cell imaging were imaged using a CorrSight spinning disk confocal microscope (Thermo Scientific) equipped with an Orca R2 CCD camera (Hamamatsu). Imaging was performed using 40x (EC Plan-Neofluar 40x/NA 1.3, Oil M27; Zeiss) or 63x (Plan-Apochromat 63x/NA 1.4, Oil M27; Zeiss) oil immersion objectives and utilizing 405, 488, 561 nm, and 633 nm lasers for excitation. Live-cell imaging was done at 5% CO_2_ supply and 37°C. Imaging of biomolecular condensates was performed on a Zeiss LSM980 confocal microscope equipped with 63x oil immersion objectives and Airyscan detection. The images were scanned at 940 × 940 pixels, unidirectionally with SR-8Y mode. Detection gain was within the range of 650 to 850. Confocal images were gain-corrected and analysed using Fiji (ImageJ) (115).

### Live-cell imaging of condensate formation

HEK293T cells were transfected with GFP-tagged Homer1, Homer2, or Homer3 using PEI as transfection reagent. After 6 hours post-transfection, the medium was replaced with phenol red–free DMEM supplemented with 10% FBS. Images were captured at 3 min intervals for approximately 5 hours using a CorrSight spinning disk confocal microscope (ThermoScientific) equipped with an Orca R2 CCD camera (Hamamatsu) and an environmental control chamber maintaining cells at 37°C and 5% CO₂ during imaging. Fluorescence excitation was achieved using a 488 nm laser for GFP, using minimal laser power and exposure times to prevent photobleaching and phototoxicity. Z-stacks using a 1 µm step size were acquired to visualise condensates throughout the cell volume. Time-lapse data was analysed using Fiji (ImageJ) (115).

### Hyperosmotic Stress Induction Using Sorbitol

For osmotic stress experiments, cells were seeded onto glass coverslips and allowed to reach 80% confluency prior to treatment. Hyperosmotic stress was induced by supplementing pre-warmed culture medium with D-sorbitol (Sigma # S1876) to a final concentration of 0.4 M for either 30 min or 60 min at 37 °C. Control cells were maintained in parallel under identical conditions without sorbitol treatment. To preserve sorbitol-induced condensates, subsequent washing and fixation steps were performed in buffers containing 0.4 M sorbitol to maintain hyperosmotic conditions. Cells were gently washed once with PBS supplemented with 0.4 M sorbitol and then fixed with 4% paraformaldehyde prepared in PBS containing 0.4 M sorbitol for 15 min at room temperature.

### Fluorescence Recovery After Photobleaching

HEK293T Cells were seeded onto Fibronectin-coated µ-Slide 8-Well Glass Bottom (Ibidi # 80827-90) at a density of 1×10^5 cells per well and transfected with mCherry and/or GFP plasmid with a total concentration of 0.26 µg of DNA with 1.3 µL of PEI as transfection reagent. FRAP Experiments were conducted 24-hours post-transfection in phenol red–free DMEM supplemented with 10% FBS. Live-cell imaging and FRAP were carried out using a 40x oil objective on Zeiss LSM 980 controlled by ZEN blue. Fluorescence excitation was achieved using 587 nm for mCherry or 488 nm for GFP laser. A circular region of interest (ROI) with a diameter of 1.5 µm was selected within a fluorescent condensate. Pre-bleach images were acquired at 0.5s intervals for 2s prior to photobleaching. Photobleaching was performed using 100% laser power for 2.45s to bleach the fluorescence within the ROI. Post-bleach recovery was monitored by acquiring images at 0.5s intervals for up to 50s, using minimal laser intensity to reduce phototoxicity and further bleaching. Fluorescence intensities were quantified using ZEN Blue by measuring the mean gray value within the bleached ROI and a reference ROI outside the bleached area to correct for overall photobleaching. Intensities were normalized to pre-bleach value and plotted as a function of time. Data were obtained from at least ten condensates per condition and plotted as mean ± SEM. Statistical analysis was performed using two-way ANOVA in GraphPad Prism.

### Image analysis

For **linescan analysis** across cell junctions and condensates, a straight line was drawn across regions of interest using the “Straight Line” tool in Fiji (ImageJ). Fluorescence intensity along the line was measured using the “Analyze → Plot Profile” function, generating a plot of pixel intensity values across the selected region. For condensate linescans, raw intensity values were normalized to the maximum intensity within each profile to obtain relative fluorescence intensity. Line graphs were generated to compare fluorescence distribution between the two fluorescence channels, representing the spatial localization of each protein across the selected structure.

To quantify the **nuclear-to-cytoplasmic** distribution of YAP, fluorescence intensity measurements were performed using ImageJ. DAPI staining was used to define nuclear regions by generating nuclear masks. The mean fluorescence intensity of the YAP signal within each nucleus was measured. The cytoplasmic intensity was calculated by subtracting the nuclear signal from the total YAP signal. The nuclear-to-cytoplasmic (n/c) ratio was obtained by dividing the nuclear intensity by the cytoplasmic intensity.

**Particle analysis** was performed in Fiji (ImageJ) to quantify condensate size and frequency under isosmotic conditions and in response to sorbitol treatment. Airyscan images (offering a lateral resolution of 100-120 nm) of cells stained for endogenous Homer1 or Homer3 were adjusted using the “Threshold” function to accurately distinguish condensates from background signal, and the selected threshold range was kept constant throughout the analysis for all conditions. Binary masks were generated, and the “Analyze Particles” function was used to measure condensate area and frequency. Size filtration was set at 0.05-0.6 µm^2^ to measure particles with a diameter of ∼250-1000 nm.

Co-localization between two proteins within condensates was quantified using **Pearson’s correlation coefficient (r)** with the Coloc2 plugin in Fiji (ImageJ). Cells were manually outlined using the Freehand Selection tool in one fluorescence channel to define the region of interest (ROI). The same ROI was then applied to the corresponding area in the second channel to ensure spatial consistency.

**Scratch areas** for wound healing assays were measured using Fiji (ImageJ). The wound region was outlined using the Freehand Selection tool, and the area was calculated using the “Measure” function under the “Analyze” menu. The remaining wound area at each time point was normalized to the initial (0-hour) scratch area to calculate the percentage of wound closure over time.

### Electron Microscopy

EGFP-APEX2 tagged Homer1, Homer2, and Homer3 constructs were transfected into 293T cells using PEI as transfection reagent. Cells were fixed 16-20 hours post-transfection using 2.5% glutaraldehyde (EM grade) in 0.1 M cacodylate buffer, pH 7.4 for 1 hour and further processed for APEX2-TEM as previously described (43, 116). Electron micrographs were captured on a Tecnai T12 TEM (ThermoFisher) operated at 120 kV using an Eagle CCD camera (4k x 4k).

### RNA extraction, cDNA synthesis and RT-qPCR

RNA was extracted using PureLink™ RNA Mini Kit (Invitrogen #12183020), following the manufacturer’s protocol. To ensure the complete removal of genomic DNA from RNA preparations, 1 µg of RNA was treated with DNase I. Specifically, 1 µL of 10X reaction buffer containing MgCl₂, 1 µL of DNase I (TURBO DNA-free™ Kit, Invitrogen #AM1907), and DEPC-treated water to a final volume of 10 µL were incubated at 37°C for 1 hour. After digestion, 1 µL of 50 mM EDTA was added at 65°C for 10 minutes. A total of 1 µL of oligo d(T)15 primer (500 µg/mL; Integrated DNA Technologies), 1 µL of random hexamers (Integrated DNA Technologies), and 1 µL of 10 mM dNTPs. This mixture was incubated at 65°C for 5 minutes, then quickly chilled on ice at 4°C for 10 minutes to allow the primers to anneal. Following annealing, 1 µL of RiboLock RNase inhibitor, 4 µL of 5X SSIV buffer, 1 µL of 100 mM DTT, and 1 µL of reverse transcriptase enzyme (SuperScript IV Reverse Transcriptase, Invitrogen™ #18090010) were added to the annealed oligo mix. The mixture was then incubated at 23°C for 10 minutes, followed by incubation at 53°C for 10 minutes to facilitate the reverse transcription reaction. Finally, the reaction was incubated at 80°C for 10 minutes to inactivate the reverse transcriptase, and the reaction was held at 4°C until further processing. A concentration of 5 ng of cDNA were used for qPCR experiment using SsoAdvanced Universal SYBR green supermix. All qPCR primers are listed in Table S1. Each qPCR reaction was prepared in a total volume of 10 µL, containing 5 µL of SYBR Green Supermix, a mix of 0.5 µL of forward and reverse primer (10 µM), 1 µL of cDNA template (5 ng), and 3.5 µL of nuclease-free water. qPCRs were performed in 96-well plates using a Bio-Rad CFX96 thermocycler. To determine relative fold changes in gene expression, ΔΔCt values were normalised to GAPDH and control conditions (MDCK wild-type cells or control siRNA-transfected cells).

### RNA sequencing

RNA sequencing (RNA-seq) of Homer triple knockout (TKO) MDCK cells was performed by NovogeneAIT using an Illumina sequencing platform. Total RNA was extracted and subjected to poly-A selection prior to library preparation. A non-stranded library preparation method was employed, and sequencing was conducted using paired-end reads with a read length of 150 base pairs for each end (2 × 150 bp), resulting in a total of 300 base pairs per fragment. Approximately 80 million reads were generated per sample, consisting of 40 million reads for read 1 and 40 million reads for read 2. NovogeneAIT provided raw sequencing data along with a comprehensive report including quality control (QC) metrics, alignment statistics, and preliminary differential expression results. Downstream analysis of RNA-seq data, including differential gene expression, was performed using R.

**Heatmaps of RNA-seq** data were generated using RStudio to visualize gene expression patterns across different experimental conditions. Raw gene expression data were normalized to eliminate technical variability and log-transformed to stabilize variance across samples. The processed data were structured as a matrix with genes as rows and samples as columns. Heatmaps were generated using the pheatmap package in R, selected for its flexibility and user-friendly interface. The pheatmap function was used to create the heatmaps, with both rows (genes) and columns (samples) hierarchically clustered based on expression similarity using default distance and linkage settings.

The top 100 upregulated and downregulated genes identified from the RNA sequencing data were further analyzed for Gene Ontology (GO) enrichment using the ShinyGO 0.8 platform (http://bioinformatics.sdstate.edu/go/). This analysis allowed for the identification of significantly enriched biological processes, molecular functions, and cellular components associated with the differentially expressed genes. Gene functions were subsequently classified into specific functional groups using the **PANTHER classification system** (https://www.pantherdb.org/), facilitating a more detailed understanding of the roles these genes play in various biological pathways.

Volcano plots were generated in R to visualize differentially expressed genes based on log₂ fold change and adjusted p-values from RNA-seq analysis. Processed differential expression results, including log₂ fold change and adjusted p-values (padj), were imported into R as a data frame. Genes were classified as significantly up- or downregulated based on defined thresholds (e.g., |log₂FC| ≥ 1 and padj < 0.05). The EnhancedVolcano package was used to create the plot. The EnhancedVolcano() function was applied, with log₂ fold change on the x-axis and –log₁₀(padj) on the y-axis. Custom colour coding and labelling were used to highlight significantly regulated genes.

### Luciferase reporter assays

TEAD (8xGTIIC) and TOPFlash luciferase reporter assays were performed using the Dual-Glo® Luciferase Assay System (Promega, #E2920) to measure the transcriptional activity of TEAD and TCF/LEF, respectively. For 24-well format experiments, cells were transfected with either 0.4 µg of 8xGTIIC firefly luciferase plasmid (Addgene, #34615) or 0.4 µg of TOPFlash firefly luciferase plasmid (Addgene, #12456), along with 0.04 µg of pRL-TK Renilla luciferase control reporter vector (Promega, #E2241), which served as an internal control for transfection efficiency. After 24–48 hours post-transfection, 100 µL of Dual-Glo® Luciferase Reagent (containing lysis buffer and substrate) was added directly to the wells containing 100 µL of culture medium. The plate was gently mixed and incubated at room temperature for 10 minutes to allow development of the firefly luciferase signal. The mixture was then transferred to a 96-well white opaque plate (Thermo Fisher, #15042), and luminescence was measured using a plate reader. Firefly luciferase activity, representing TEAD or TCF/LEF transcriptional activity, was detected at 560 nm emission. Next, 100 µL of Dual-Glo® Stop & Glo® Reagent was added to each well to simultaneously quench the firefly signal and activate Renilla luciferase. Renilla luminescence, which serves as a normalization control, was measured immediately afterward, at 480 nm. Firefly luciferase values were normalized to corresponding Renilla luciferase values to account for variability in transfection efficiency. Final results were expressed as the ratio of firefly to Renilla luminescence and further normalized to wild-type or control siRNA conditions to provide a relative measure of transcriptional activity.

### Quantifications and Statistical Analysis

All statistical analyses were conducted using two-tailed paired Student’s t-test analysis in GraphPad Prism, except for qPCR analysis, which was performed using a Two-way ANOVA with Tukey’s multiple comparison tests. FRAP statistical analysis was performed using Two-way ANOVA. Significance of data is represented as ns, not significant, * P ≤ 0.05, ** P ≤ 0.01, *** P ≤ 0.001, **** P ≤ 0.0001. Error bars in all figures represent mean +/- SEM.

## Supporting information

Resources Table

## Acknowledgements

We are grateful to Boon Chuan Low and Darren Wong Chen Pei for reagents and plasmids and Jeanette Leong, Amirul Sufyan Bin Mohamed Salleh, and Shanmugavelu Balasubramaniam Ravendrakum for their contribution to this work during their Final Year Projects. This project is supported by the Ministry of Education, Singapore, under its Academic Research Fund Tier 2 (MOE-T2EP30121-0019) and Tier 1 (RG31/23).

## Author contributions

SMJMY generated and characterized Homer KO cells, performed qPCR, reporter assays, biochemical and microscopy experiments, and analysed the data; LJW generated FRYL KO cells and performed stainings on mouse kidney sections; YC generated PATJ KO cells and performed qPCR and reporter assays; BH performed the APEX2-TEM analysis of Homer condensates; AL obtained funding, conceived the project, supervised SMJMY, LJW, YC and BH, and wrote the manuscript.

