## Supplemental Figures for "Homer condensates orchestrate YAP-Wnt signaling crosstalk downstream of the Crumbs polarity complex"

### Supplemental Figures and files:

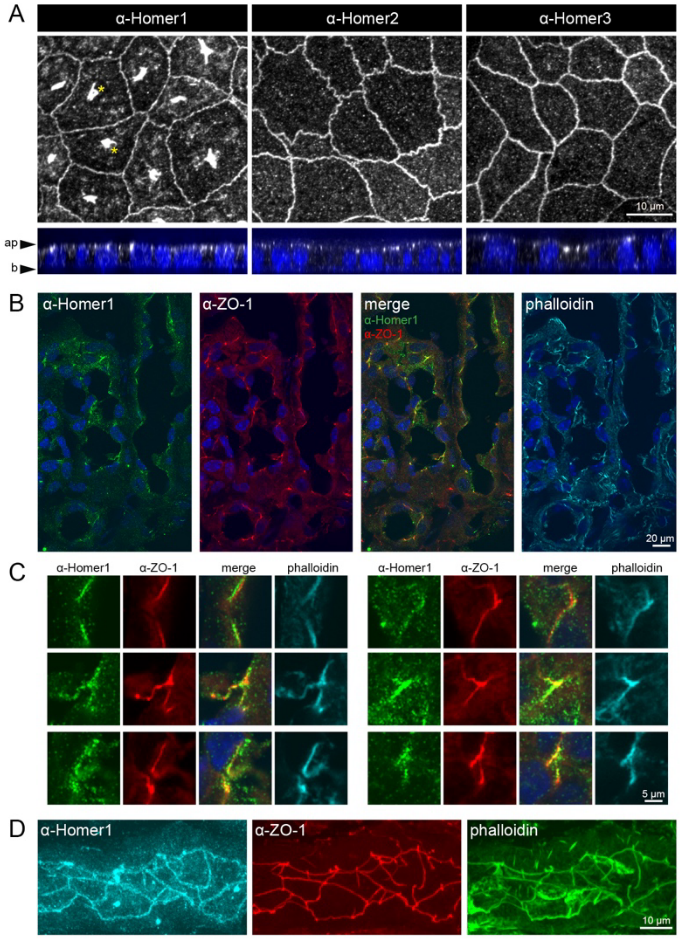


**Figure S1: Homers localise to apical cell junctions in renal epithelial cells**
(A) Localisation of Homers in MDCK-II cells. Cells were grown to confluency on Transwell filters, fixed and stained with Homer1, Homer2 or Homer3 antibodies. XY (en face) and XZ (side view) projections of confocal z-stacks are shown. Note that anti-Homer1 antibodies produce a non-specific staining of unknown origin in the cytoplasm (yellow asterisks). (B-D) Localisation of Homer1 in mouse kidney cryosections. Sections were stained with antibodies against Homer1 and ZO-1 and counterstained with phalloidin and DAPI. (B) Cross-section through renal tubules. (C) Magnified views of apical cell junctions based on the images shown in (B). (D) En face sections through apical cell junctions in renal epithelial cells.

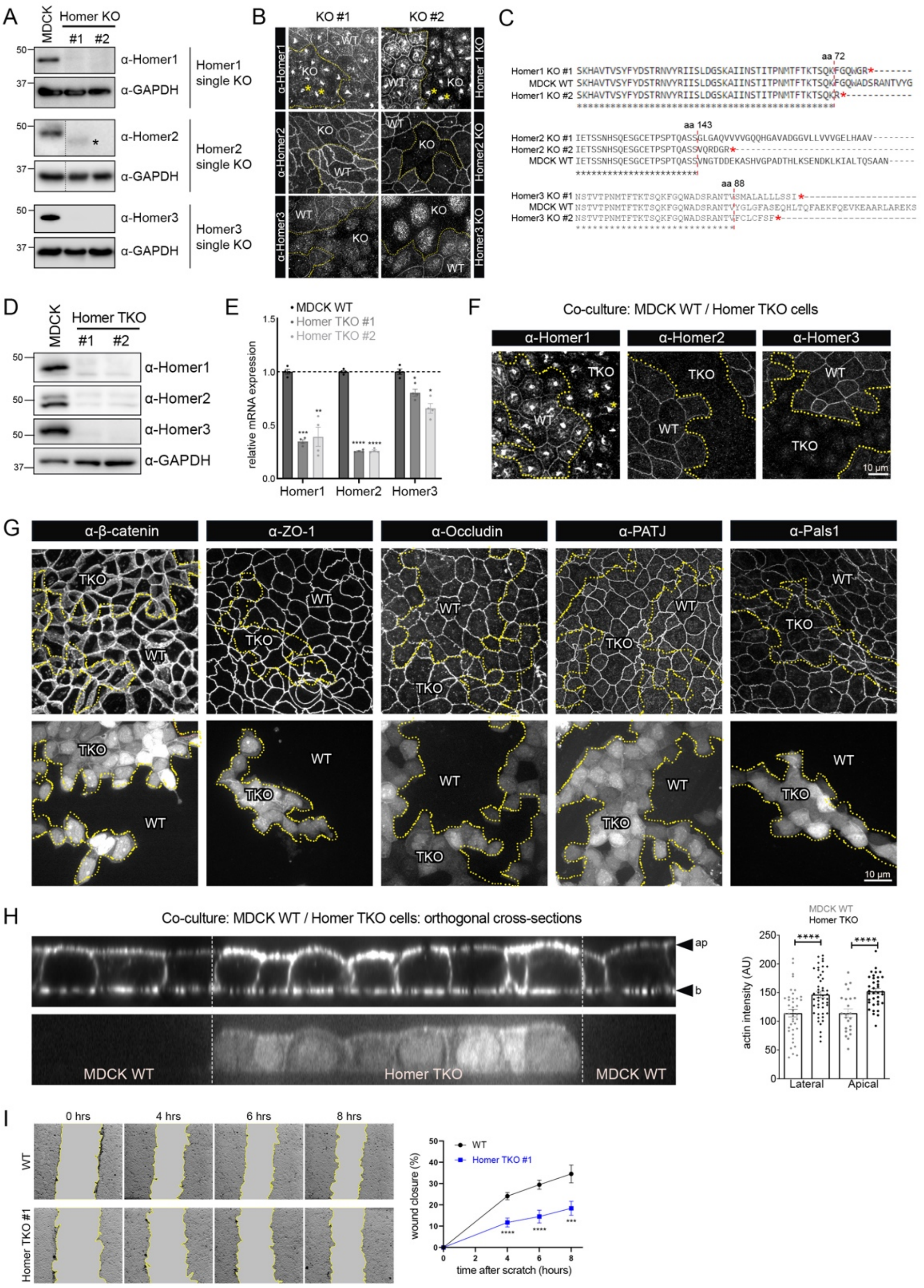


**Figure S2: Generation and characterisation of Homer KO cells**

(A) WB analysis of single Homer1, Homer2, and Homer3 knockout (KO) MDCK cell lines. Note that a truncated version of Homer2 protein is detected in KO clone #1 (asterisk). (B) MDCK WT cells were co-cultured with single Homer1, Homer2 or Homer3 KO cells on Transwell filter and stained for the corresponding Homer protein. Dotted lines represent the boundary between the single KO cells and MDCK WT cells. Note that anti-Homer1 antibodies produce a non-specific, non-junctional staining that persists in Homer1 KO cells (yellow asterisks). Note also that Homer2 protein is detected but reduced in Homer2 KO clone #1. (C) Amino acid sequence alignment of single Homer1, Homer2, and Homer3 KO cells compared to MDCK WT at their respective loci. Red asterisks (*) indicate premature stop codons. The red line marks the point at which the sequence between KO and WT genomic DNA diverges. (D) Validation of Homer TKO cells. WB analysis of Homer TKO and MDCK WT cell lysates probed for all three Homer proteins. (E) qPCR analysis of Homer1, Homer2 and Homer3 mRNA expression in Homer TKO cell lines (n=2). (F) Homer TKO and MDCK WT cells were co-cultured on Transwell filters for 10-14 days, fixed and stained for Homer1, Homer2 or Homer3. The dotted lines represent the boundaries between the two cell populations. (G) Homers are not required for the assembly or maintenance of the apical junctional complex. MDCK WT and Homer TKO (GFP positive) cells were co-cultured on Transwell filters, fixed and stained for junctional markers β-catenin, ZO-1, Occludin, PatJ, and Pals1. The dotted lines delineate the boundaries between WT and Homer TKO cell populations. (H) Loss of Homers increases cortical actin levels. MDCK WT and Homer TKO (GFP positive) cells were co-cultured on Transwell filters, fixed and stained with phalloidin. Orthogonal cross-sections through the monolayer and quantification of phalloidin intensity at the apical and lateral membranes are shown (n=2 independent co-cultures). Data is presented as mean ± SEM. Statistical significance is ****$P \leq 0.0001$. (I) Scratch wound assay of confluent WT MDCK cells and Homer TKO cells. . Wound areas were quantified using ImageJ and normalised to the 0-hour time point (n=3). Data is presented as mean ± SEM. Statistical analysis was performed using two-way ANOVA. * $P \leq 0.05$, ** $P \leq 0.01$, *** $P \leq 0.001$, **** $P \leq 0.0001$.

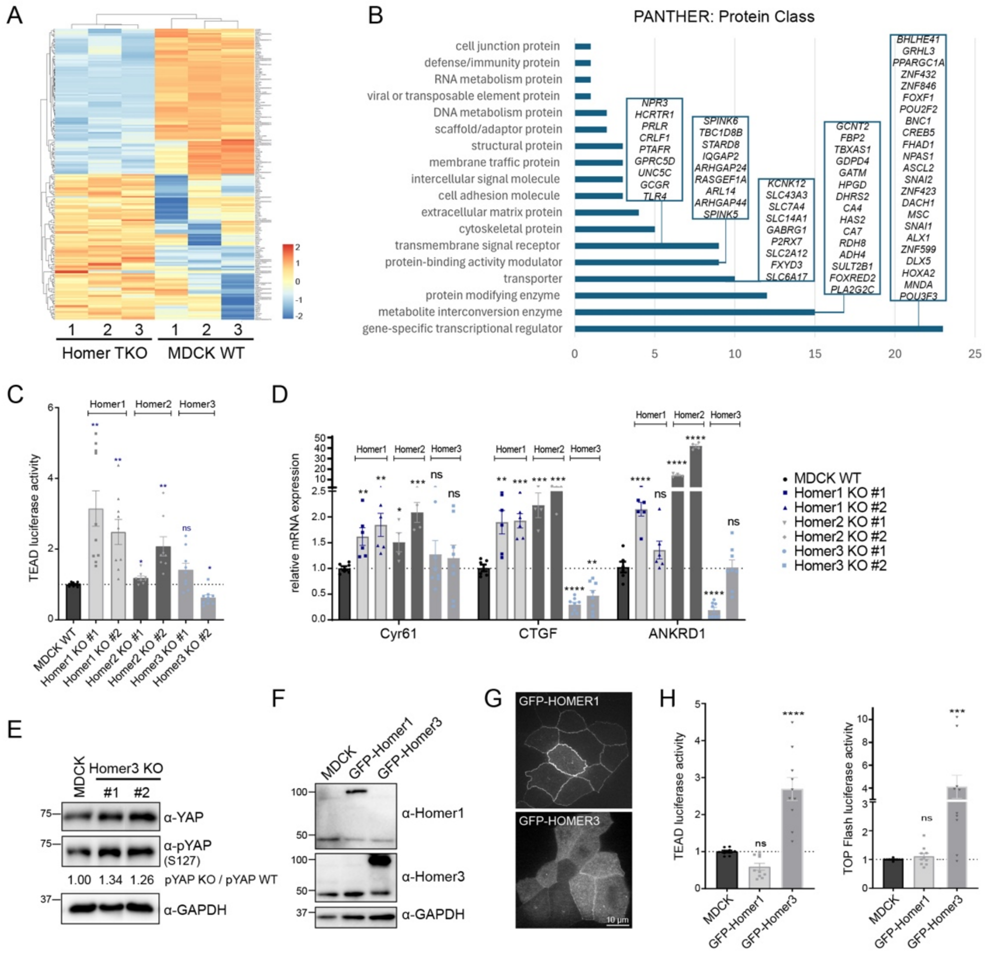


**Figure S3: Homers differentially regulate YAP/TEAD activity in MDCK cells**

(A) Heatmap showing the top 100 upregulated and top 100 downregulated genes in Homer TKO MDCK cells compared to MDCK WT cells, as identified by RNA-seq analysis (n=3). Colour intensity represents relative expression levels, with red indicating upregulation and blue indicating downregulation. See Table S2 for a complete list of all differentially expressed genes along with their log2 fold changes and raw RNA-seq output. (B) The top 100 upregulated and top 100 downregulated genes were classified by PANTHER: Protein Class. Genes in the classification groups are listed in boxes. (C) TEAD luciferase activity in single Homer1, Homer2 and Homer3 KO cells normalised to MDCK WT cells (n=4). (D) qPCR analysis of YAP target gene expression in single Homer1, Homer2 and Homer3 KO cells normalised to MDCK WT cells (n=3 or 4). (E) WB analysis of MDCK WT and Homer3 KO cell lysates probed for YAP and pYAP (S127). The pYAP/YAP ratio was quantified by densitometric analysis (n=3). (F) WB analysis of MDCK WT cells and MDCK cells stably transfected with GFP-Homer1 or GFP-Homer3. (G) Representative confocal micrographs of MDCK cells stably transfected with GFP-Homer1 or GFP-Homer3 grown on Transwell filter for 10-14 days. (H) TEAD and TOPFlash luciferase assays of MDCK WT cells and MDCK cells stably transfected with GFP-Homer1 or GFP-Homer3 (n=5). Data is presented as mean ± SEM. Statistical significance is indicated as follows: *P ≤ 0.05; **P ≤ 0.01; ***P ≤ 0.001; ****P ≤ 0.0001.

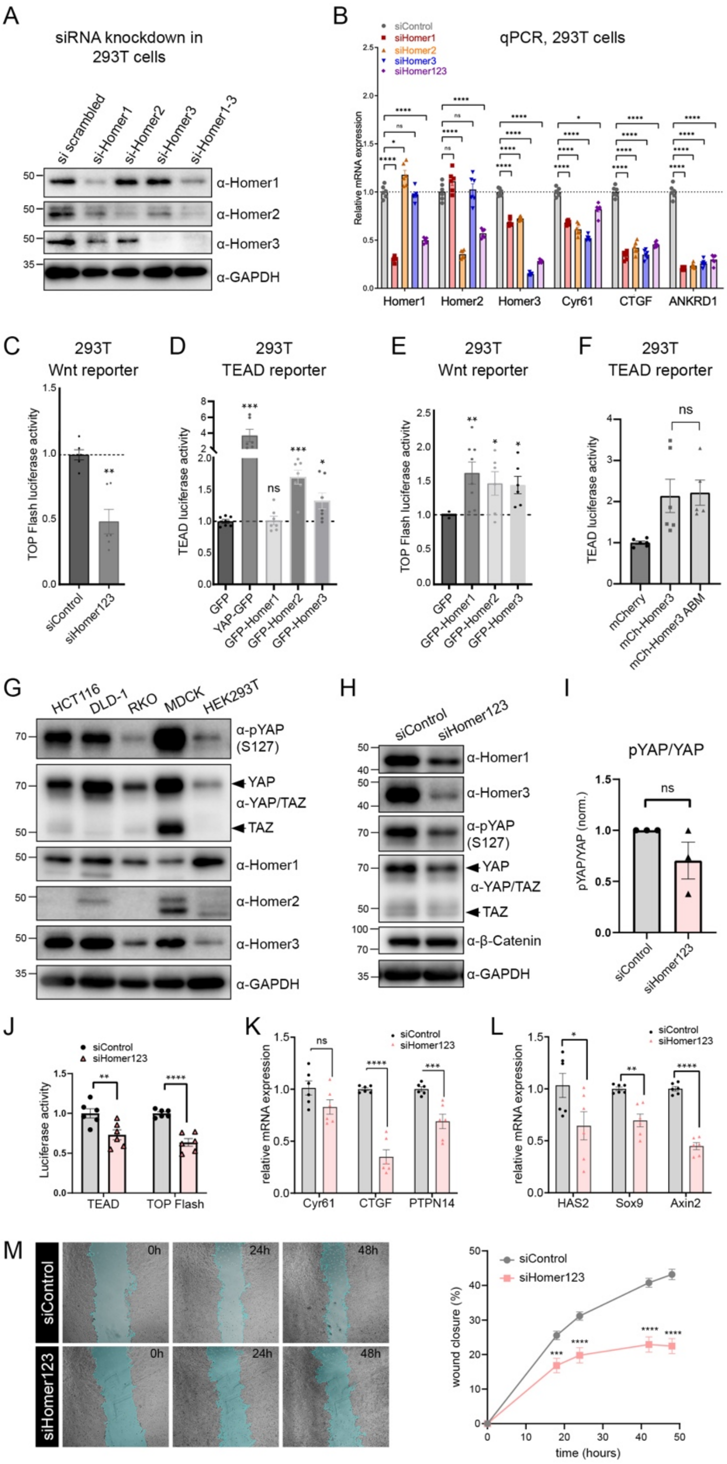


**Figure S4: Homers promote YAP/TEAD and Wnt pathway activity in HEK293T and HCT116 cells**

(A and B) WB and qPCR analysis of 293T cells transfected with a total of 100 nM siRNA targeting either Homer1, Homer2, or Homer3, or all three Homers simultaneously (n=3). (C) TOPFlash luciferase activity of 293T cells transfected with a total of 100 nM of control siRNA or a pool of siRNAs targeting all three Homers (n=3). (D) TEAD luciferase assay of 293T cells transfected with GFP or GFP-tagged YAP, Homer1, Homer2 or Homer3 (n=3). (E) TOPFlash luciferase activity in 293T cells transfected with GFP or GFP-tagged Homer1, Homer2, or Homer3 (n=3). (F) TEAD luciferase assay of 293T cells transfected with mCherry-Homer3 or a corresponding actin binding mutant (ABM, R45E, R49E, R84E) of Homer3 (n=3). (G) WB analysis of three colorectal cancer cell lines (HCT116, DLD-1, and RKO) and two non-cancerous cell lines (MDCK-II and HEK293T), probed for the expression of YAP, phosphorylated YAP (S127), Homer1, Homer2, and Homer3. (H) WB analysis of HCT116 cells treated with control siRNA or a pool of siRNAs targeting all three Homers. Cell lysates were collected 48 hours post-transfection and probed for Homer1, Homer3, YAP, pYAP (S127), and β-catenin. (I) Quantification of the pYAP/YAP ratio based on Western blotting in (H) (n=3). (J) TEAD and TOPFlash reporter assays in HCT116 cells transfected with control siRNA or a pool of siRNAs targeting all three Homers (n=3). (K and L) qPCR analysis of YAP target genes (K) and WNT-associated genes (L) in siRNA-transfected HCT116 cells. (M) Scratch wound assay of siRNA transfected HCT116 cells. Following siRNA transfection, cells were serum-starved for 6 hours prior to scratch induction and maintained in low-serum media. Wound areas were quantified using ImageJ and normalised to the 0-hour time point (n=3). Data is presented as mean ± SEM. Statistical analysis was performed using two-way ANOVA. * $P \leq 0.05$, ** $P \leq 0.01$, *** $P \leq 0.001$, **** $P \leq 0.0001$. All Luciferase assays were statistically analysed using a paired Student's t-test. All qPCR data were analysed using Two-Way ANOVA Tukey multiple comparison test. Data is presented as mean ± SEM. *$P \leq 0.05$; **$P \leq 0.01$; ***$P \leq 0.001$; ****$P \leq 0.0001$.

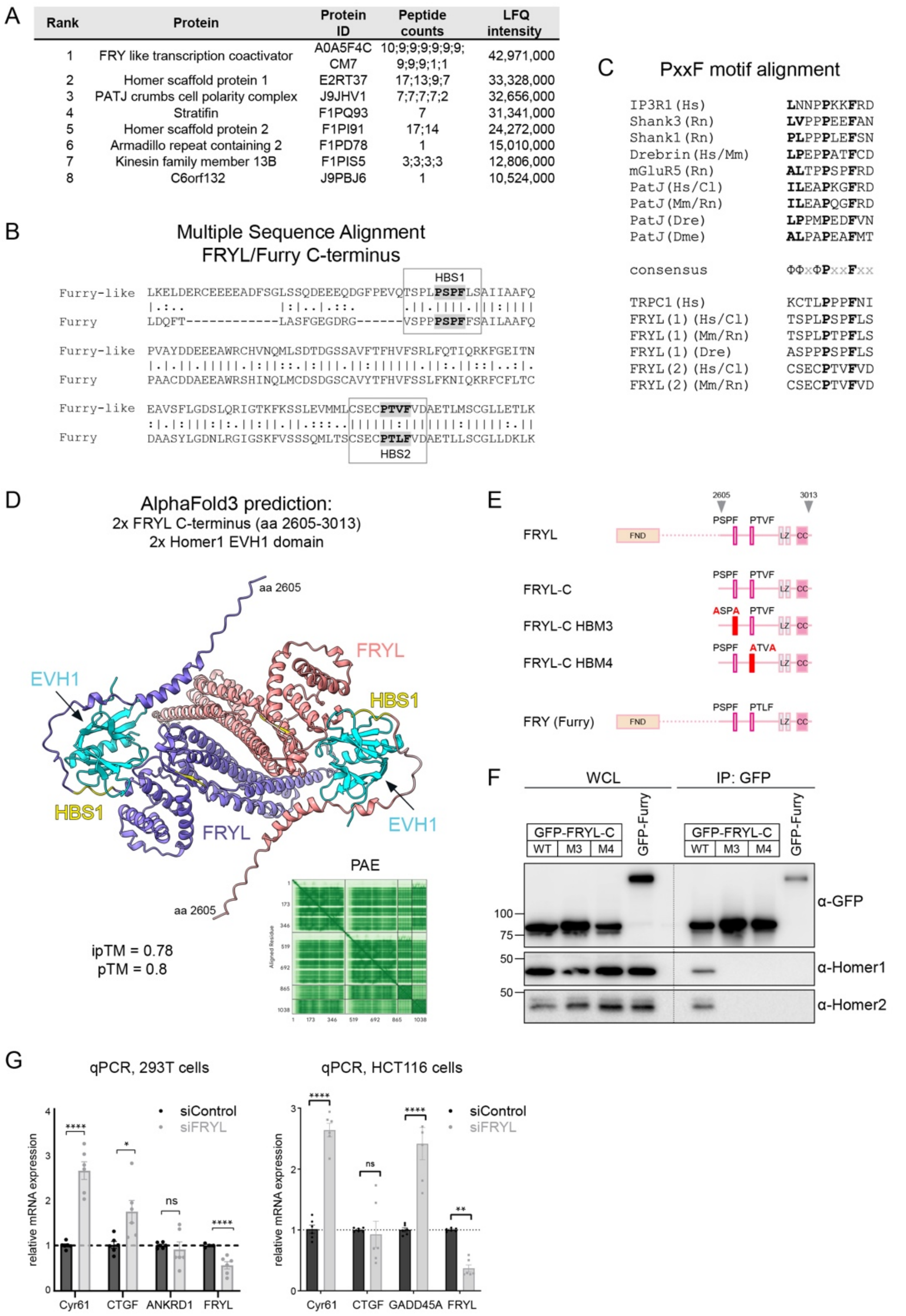


**Figure S5: FRYL interacts with Homer proteins via two PxxF motifs**

(A) IP-MS data of GFP-Homer3 stably overexpressed in MDCK cells. The table shows the top-ranked proteins based on peptide count identified in the MS results. (B) Amino acid sequence alignment of the C-terminus of human FRYL and Furry. The two PxxF motifs (Homer binding sites (HBS)) in FRYL are highlighted. (C) Amino acid sequence alignment of PxxF motifs from known Homer EVH1 domain ligands and those identified in PatJ and FRYL. Mm: *Mus musculus*, Rn: *Rattus norvegicus*, Cl: *Canis lupus familiaris*, Hs: *Homo sapiens,* Dre*: Danio rerio*, Dme: *Drosophila melanogaster*. (D) AlphaFold3 prediction of the FRYL C-terminus dimer in complex with two Homer1 EVH1 domains. The HBS1 and HBS2 are highlighted in yellow. HBS1 is predicted to interact with the hydrophobic groove of the EVH1 domains, whilst HBS2 is buried. The Predicted Alignment Error (PAE) plot and the ipTM and pTM values are shown. (E) Cartoon depicting the two Homer binding mutants (HBM) M3 and

M4 generated in the context of the FRYL C-terminus (FRYL-C). (F) IP of GFP-Furry, GFP-FRYL-C, and the two GFP-FRYL-C HBMs M3 and M4 from 293T cell lysates. Note that mutation of either of the two PxxF motifs abolishes the interaction with Homers. Furry does not interact with Homers although it contains two PxxF motifs that are similar in primary sequence to those found in FRYL (B). (G) qPCR analysis of canonical YAP target genes in 293T cells (left) and HCT116 cells (right) transfected with control or FRYL siRNA. qPCR data were analysed using Two-Way ANOVA (n=6). Data is presented as mean ± SEM. * $P \leq 0.05$, ** $P \leq 0.01$, *** $P \leq 0.001$, **** $P \leq 0.0001$.

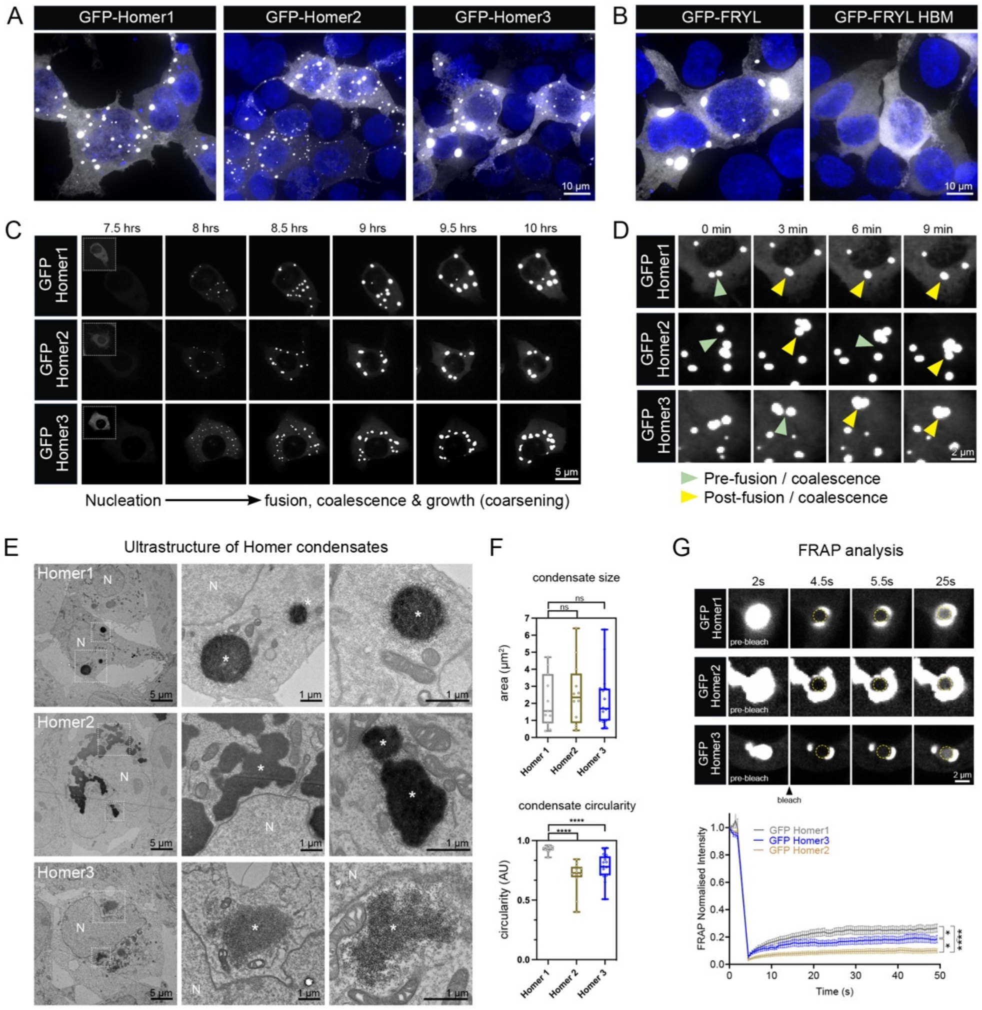


**Figure S6: Characterisation of biomolecular condensates induced by transient overexpression of GFP-Homers in 293T cells**

(A) Confocal micrographs of 293T cells transiently transfected with GFP-Homer1, GFP-Homer2 or GFP-Homer3. (B) Confocal micrographs of 293T cells transiently transfected with WT GFP-FRYL or GFP-FRYL HBM M2. (C) 293T cells were transfected with GFP-tagged Homer constructs and imaged live 6 hours post-transfection using spinning disk microscopy. Representative time frames illustrating the nucleation, growth and maturation of Homer condensates are shown. The gain in the insets shown at 7.5 hrs was increased to show diffuse cytoplasmic localisation. (D) Live cell imaging illustrating the fusion and coalescence of condensates formed by GFP-tagged Homer1, Homer2, and Homer3. Green arrowheads indicate the prospective site of fusion, yellow arrowheads mark the post-fusion structures. (E) Representative transmission electron micrographs of 293T cells transfected with APEX2-EGFP tagged Homer1, Homer2 or Homer3. EM contrast of Homer condensates was enhanced through APEX2 labeling. Boxed areas are magnified. Note the distinct ultrastructure of Homer condensates. N=nucleus. (F) Quantification of condensate size and circularity based on TEM. ****$P \leq 0.0001$ (One-Way ANOVA, n=12-16 condensates). (G) FRAP analysis of GFP-Homer1, GFP-Homer2 or GFP-Homer3 transiently transfected into 293T cells. Data is presented as mean ± SEM. Statistical analysis was performed using two-way ANOVA. ** $P \leq 0.01$.

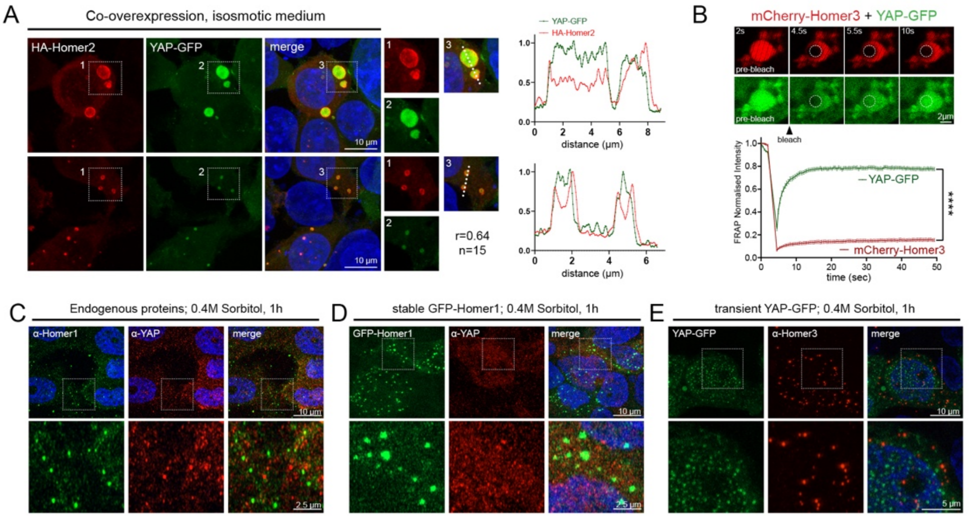


**Figure S7: YAP is excluded from Homer condensates formed at or close to endogenous expression levels**
(A) 293T cells transiently co-transfected with HA-Homer2 and YAP-GFP. Examples of large (top) and small (bottom) condensates are shown. (B) FRAP analysis of 293T cells co-transfected with mCherry-Homer3 and YAP-GFP (n=31). FRAP data is presented as mean ± SEM. Statistical analysis was performed using two-way ANOVA. ** $P \leq 0.01$, *** $P \leq 0.001$, **** $P \leq 0.0001$. (C) Wild-type 293T cells were exposed to 0.4M sorbitol for 1h, fixed and stained with antibodies against Homer1 and YAP. (D) 293T cells stably transfected with GFP-Homer1 were exposed to 0.4M sorbitol for 1h, fixed and stained with anti-YAP antibodies. (E) 293T cells transiently transfected with YAP-GFP were exposed to 0.4M sorbitol for 1h, fixed and stained with anti-Homer1 antibodies. Images in C-E were acquired in Airyscan mode.

**Supplemental Tables:**

**Table S1: qPCR primers and oligonucleotides**
**Table S2: RNA sequencing data.** Differentially expressed genes confirmed by qPCR are highlighted.

**Supplemental Movies:**

**Movie S1:** Formation of GFP-Homer1 condensates in response to osmotic stress
**Movie S2:** Fusion of GFP-Homer1 condensates in sorbitol-treated cells
**Movie S3:** GFP-FRYL and mCherry-Homer3 form droplet-like condensates in live cells
**Movie S4:** mCherry-PATJ and GFP-Homer1 rapidly coalesce into interconnected networks upon osmotic stress induction
**Movie S5:** GFP-PATJ PDZ1-8 and mCherry-Homer3 form droplet-like condensates in live cells
